# Combination of Polymeric Micelle Formulation of TGFβ Receptor Inhibitors and Paclitaxel Produce Consistent Response Across Different Mouse Models of TNBC

**DOI:** 10.1101/2023.06.14.544381

**Authors:** Natasha Vinod, Duhyeong Hwang, Sloane Christian Fussell, Tyler Cannon Owens, Olaoluwa Christopher Tofade, Sage Copling, Jacob D. Ramsey, Patrick D. Rädler, Hannah M. Atkins, Eric E. Livingston, J. Ashley Ezzell, Marina Sokolsky- Papkov, Hong Yuan, Charles M. Perou, Alexander V. Kabanov

## Abstract

Triple-negative breast cancer (TNBC) is notoriously difficult to treat due to the lack of targetable receptors and sometimes poor response to chemotherapy. The transforming growth factor-beta (TGFβ) family of proteins and their receptors (TGFR) are highly expressed in TNBC and implicated in chemotherapy-induced cancer stemness. Here we evaluated combination treatments using experimental TGFR inhibitors (TGFβi), SB525334 (SB), and LY2109761 (LY) with Paclitaxel (PTX) chemotherapy. These TGFβi target TGFR-I (SB) or both TGFR-I&II (LY). Due to the poor water solubility of these drugs, we incorporated each of them in poly(2-oxazoline) (POx) high-capacity polymeric micelles (SB-POx and LY-POx). We assessed their anti-cancer effect as single agents and in combination with micellar Paclitaxel (PTX-POx) using multiple immunocompetent TNBC mouse models that mimic human subtypes (4T1, T11-Apobec and T11-UV). While either TGFβi or PTX showed a differential effect in each model as single agents, the combinations were consistently effective against all three models. Genetic profiling of the tumors revealed differences in the expression levels of genes associated with TGFβ, EMT, TLR-4, and Bcl2 signaling, alluding to the susceptibility to specific gene signatures to the treatment. Taken together, our study suggests that TGFβi and PTX combination therapy using high-capacity POx micelle delivery provides a robust anti-tumor response in multiple TNBC subtype mouse models.

**Translational Impact Statement:** Paclitaxel is a widely used chemotherapy in breast cancer. However, response to single-agent chemotherapy is short-lived in a metastatic setting. This study shows the broad applicability of the therapeutic combination of TGFβ inhibitors with Paclitaxel across different TNBC subtypes.

## Introduction

According to the Global Cancer Statistics, there were 19 million new cancer cases and 10 million cancer deaths estimated worldwide in 2020 [1]. Female breast cancer accounted for 11.7% of all cancer types (2.3 million cases), making it the most common type of cancer in 2020 and the leading cause of cancer death in women. Triple-negative breast cancers (TNBC), characterized by the lack (less than 1%) of receptors for estrogen, progesterone, and lack of human epidermal growth factor 2 (HER 2) overexpression, have the worst prognosis of all breast cancers. The analysis of gene expression patterns in different tumors has uncovered several main subtypes of breast cancer. These include basal-like, luminal A, luminal B, HER-2 enriched, and Claudin-low [2]. Unlike the luminal subtype, the basal-like subtype is associated with poor prognosis and is distinguished from luminal tumors based on the expression of cytokeratins. The majority of TNBCs have a basal epithelial-like phenotype and stain positive for cytokeratins 5/6 and 17 [3]. The Claudin-low TNBC subtype, discovered in 2007 by Herschkowitz et al., is classified by the low expression of Claudin 3, 4, and 7 and other tight junction proteins such as E-cadherin [2]. In addition to the genomic and histopathological heterogeneity, TNBC is also heterogeneous at the transcriptomic level and has been further divided into 7 clusters on the basis of biological pathway activity [4].

Due to the lack of targetable receptors and sometimes poor response to immunotherapy using immune checkpoint inhibitors (ICI), TNBC has been notoriously difficult to treat. Until recently, cytotoxic chemotherapy has been the mainstay of TNBC treatment [5]. However, although TNBC responds to chemotherapy to some extent, the median overall survival with single-agent chemotherapy is only 17.5 months in the metastatic setting [6]. More recently developed targeted therapeutics, such as the Poly (ADP-ribose) polymerase (PARP) inhibitors, Olaparib and Talazoparib, have shown promising outcomes and were approved in patients with BRCA mutations [7]. However, around 40% to 70% of the patients are likely to develop resistance to these therapies [8]. Moreover, the significant level of heterogeneity of TNBC makes it practically challenging to stratify patients based on specific genomic and molecular signatures and identify candidates for targeted therapies. Besides, the heterogeneity of TNBC grows over time, resulting in the emergence of drug-resistant cancer stem-like cells [9]. The high relapse rates and poor prognosis of TNBC have fueled efforts to find new treatments that could be effective across different TNBC subtypes.

Combining chemotherapy with ICI has shown promise for treating TNBC. Clinical trial results showed that pembrolizumab (anti-PD1) had an improved response rate when paired with chemotherapy (nab-paclitaxel or paclitaxel, or gemcitabine plus carboplatin) than when used alone. These results granted accelerated Food and Drug Administration (FDA) approval of pembrolizumab and chemo-immunotherapy combinations for PDL1-positive TNBCs [10]. Atezolizumab (anti-PD-L1) in combination with nab-paclitaxel also received an accelerated FDA approval for the treatment of patients with unresectable locally advanced or metastatic PD-L1 positive TNBC based on the initial Phase 3 clinical trial [11]; however, data from the subsequent additional Phase 3 trial showed that the combination of atezolizumab and paclitaxel did not improve progression-free survival compared to paclitaxel alone (https://clinicaltrials.gov/ct2/show/NCT03125902). Subsequently, the combination of atezolizumab and chemotherapy was withdrawn (https://www.gene.com/media/press-releases/14927/2021-08-27/genentech-provides-update-on-tecentriq-u). Given this development, as well as the fact that ICI treatment is associated with severe immune-related adverse events, the need for newer therapeutic strategies that can blend well with chemotherapy while causing fewer side effects remains important.

The transforming growth factor-beta (TGFβ) is a superfamily of pleiotropic cytokines that regulate various biological processes, including cell growth, differentiation, immune regulation, and apoptosis [12]. TGFβ signaling negatively regulates the inflammatory response of immune cells, thereby promoting tumor growth [13]. Cancer cells exploit TGFβ signaling to promote EMT, immune tolerance, and myofibroblast differentiation [14]. TGFβ proteins have recently been found to play a key role in regulating stemness in breast cancer cells, suggesting that it could be a crucial target in TNBC [15]. Biopsies from breast cancer patients who received chemotherapy showed an increase in TGFβ and cancer stem cell (CSC) markers compared to biopsies obtained before the treatment [16]. The expression of CSC gene signatures (CD44 and ALDH1A1) in human breast cancer cells with high autocrine TGFβ signaling can be downregulated by a TGFβ inhibitor (TGFβi) that targets TGFβ receptor-I (TGFR-I) [16]. These findings support the rationale for combining chemotherapy with inhibitors of TGFβ receptors in TNBC.

Here we evaluated two small-molecule TGFβi, SB525334 (SB) and LY2109761 (LY), that target TGFR-I or both TGFR-I and TGFβ receptor-II (TGFR-II), respectively. Since these drugs are water-insoluble, we incorporated them in the polymeric micelles (PMs) of poly(2-oxazoline) (POx) block copolymer to enable parenteral delivery of poorly soluble agents [17]. We used the micellar drugs (SB-POx and LY-POx, respectively) as single treatments and in combination with paclitaxel (PTX), which was also incorporated in POx PMs (PTX-POx). For the therapeutic assessment, we employed syngeneic TNBC models that are representative of a hybrid basal-like and Claudin-low phenotype (4T1) and Claudin-low phenotype (T11-Apobec and T11-UV).

## Materials and Methods

### Materials

The triblock copolymer P[MeOx_36_-b-BuOx_21_-b-MeOx_35_]-piperadine (POx) [M_w_/M_n_ (number-average molecular weight/weight-average molecular weight) = 1.04] was synthesized by ring-opening polymerization of 2-oxazolines as described previously [18]. The block lengths were determined using ^1^H nuclear magnetic resonance (NMR) spectroscopy on a Bruker Avance III 400-MHz spectrometer, and the NMR spectrum was analyzed using the MestReNova software (11.0). The molecular weight distribution of POx was obtained by gel permeation chromatography on a Viscotek VE2001 solvent sampling module. TGFβi, LY, and SB, were purchased from Adooq BioScience and APExBIO, respectively. PTX was purchased from LC laboratories. The remaining reagents were purchased from Sigma-Aldrich or Fisher Scientific unless specified otherwise.

### Cell Culture

NIH-3T3 mouse embryonic fibroblasts and 4T1 mammary carcinoma cell lines were obtained from the Tissue Culture Facility at UNC Lineberger Comprehensive Cancer Center (LCCC). The NIH-3T3 cells were cultured in DMEM (Gibco) medium supplemented with 10% fetal calf serum and 1% penicillin and streptomycin at 37^0^C in a humidified atmosphere containing 5% CO_2_. The 4T1 TNBC cells were obtained from UNC LCCC Tissue Culture Facility and were cultured in the same way, except that the basal media was RPMI (Gibco), and fetal bovine serum was used instead of fetal calf serum. The T11-Apobec and T11-UV TNBC cells were provided by Dr. Charles M. Perou (LCCC) and cultured in RPMI medium (Gibco 11965-092) supplemented with 5% FBS, 1% penicillin-streptomycin and puromycin.

### Preparation and characterization of POx micelles

The POx micelles of SB, LY, PTX, and LY/PTX (4/8), SB/PTX (5.2/8), LY/PTX (5.2/8) were prepared using the thin-film hydration as recently described [17]. Both the drugs and polymer were dissolved in ethanol, which served as a common solvent. The details of the evaporation and hydration conditions for each formulation are mentioned in **Table S1**. The drug loading into micelles was measured using reversed-phase high-pressure liquid chromatography on an Agilent 1200 HPLC system under isocratic elution mode with a mobile phase of acetonitrile/water 70/30 (v/v) containing 0.1% trifluoroacetic acid operated at a flow rate of 1 ml/min. A Nucleosil C18, 5um particle size column [*L* × inner diameter (ID) 25 cm by 4.6 mm] was used, and the column temperature was set to 40^0^C. The micelle samples were diluted 50-fold in acetonitrile/water 50/50 (v/v), and an injection volume of 10 uL was used. The drug loading capacity of the POx formulations was measured as described previously. Further, the size distribution of the formulations was determined using dynamic light scattering (DLS) on a Zetasizer Nano ZS (Malvern Instruments Ltd., UK). The samples were diluted 10-fold in saline, and three measurements were recorded for each sample at 25^0^C. POx micelles were also characterized by transmission electron microscopy (TEM). The drug-loaded micelles were diluted 10-fold in distilled water to a final POx concentration of 2 mg/mL, dropped onto a TEM grid (Ted Pella Inc, Carbon Type-B, 200 mesh, Copper), allowed to dry, and then stained for 2 min with 1% uranyl acetate before TEM imaging. The Thermo Scientific Talos F200X scanning/transmission electron microscope (accelerating voltage of 200kV) was used for image acquisition.

### *In vitro* TGFβ inhibition

The NIH-3T3 cells were seeded in a 12-well plate and grown overnight. The media was replaced after 24h with fresh media containing 2.5 ng/ml TGFβ1 ligand (R&D Systems), with or without 5µM TGFβi (SB-POx, unformulated SB in DMSO (SB), LY-POx, unformulated LY in DMSO (LY)). Saline or vehicle (POx only or DMSO) were used as controls. Two hours after treatment, the cells were washed with PBS and lysed with RIPA buffer (EMD Millipore) containing phosphatase and protease inhibitor cocktail (Sigma) and frozen overnight. The samples were thawed the next day, subjected to intermittent sonication, and centrifuged at 17,000 g for 10 min at 4^0^C. The supernatant containing the whole cell lysate was collected for further analysis. The total protein concentration in the samples was measured using the Pierce BCA Protein Assay kit (Thermo Scientific). ProteinSimple Wes, a capillary-based western blotting system, was used to assess the expression of phosphorylated Smad2/3 (p-Smad2/3). The housekeeping protein was β-actin. The antibodies were titrated to determine the optimal concentration with the least non-specific binding. **Table S2** contains information on the primary antibody clone, concentration, and lysate concentration. The anti-rabbit secondary antibody and all other reagents were obtained from ProteinSimple. The NIH-3T3 cells were treated with TGFβ1 with or without TGFβi in the same manner as in the western blot experiment. Cells not stimulated with TGFβ1 were used as unstimulated control. One hour after treatment, the cells were washed with PBS, detached with 100uL of TrypLE Express (Gibco) treatment for 2 min, and neutralized with 1 mL medium. The cell suspension was then added to 10 volumes (10mL) of 1x Lyse/Fix buffer (BD Phosphoflow) prewarmed to 37^0^C and incubated for 12 min at that temperature. The cell suspension was centrifuged at 600g for 7 min, and the supernatant was removed, leaving about 50uL residual volume. The cell pellet was washed with 10mL of BD stain buffer to remove traces of Lyse/Fix buffer and centrifuged again. After that, the cells were permeabilized by resuspending the cell pellet in 1mL of pre-chilled Perm Buffer III (BD Phosphoflow) and incubated on ice for 30 min. The cells were pelleted and washed three times with the BD stain buffer and resuspended at a final concentration of 10M cells/mL with BD stain buffer before adding Fc block antibody (TruStain; diluted 50-fold). Following a 15 min incubation, PE anti-p-Smad2/3 antibody (see **Table S2** for antibody information) was directly added to the cell suspension, and the cells were incubated for 60 min at RT (protected from light). A small percentage of cells were set aside as unstained control (USC) and did not receive the antibody stain. Following antibody staining, the cells were pelleted and washed twice with BD stain buffer, resuspended with 200uL BD stain buffer, and mixed thoroughly before being analyzed on Attune NxT flow cytometer (Thermo Fisher).

### Cytotoxicity

Eight thousand 4T1 cells were seeded per well in 96-well plates and allowed to grow overnight. The following day, SB-POx, LY-POx, and PTX-POx were serially diluted in the cell culture medium at 5-fold increments, and nine different concentrations of each drug ranging from 0.256 nM to 100 μM were tested on cells. Wells that only received the media served as control. Twenty-four hours following treatment, the cell viability was determined using the Dojindo Cell Counting Kit-8 (CCK-8) using the manufacturer’s instructions. The half-maximal inhibitory concentration (IC_50_) was calculated using the GraphPad Prism 9 software.

### Tumor Inhibition

Animal studies were conducted in accordance with the University of North Carolina at Chapel Hill Institutional Animal Care and Use Committee guidelines. One million 4T1 cells (in 100uL of 1:1 mix of Hank’s balanced salt solution and BD Matrigel) were injected into the 4^th^ left inguinal mammary fat pad of 8-week-old female BALB/c mice. For the T11-Apobec and T11-UV TNBC models, 0.1 million cells (in 100 µL of 1:1 mix of PBS and Cultrex Basement Membrane Extract (Type 3, R&D systems)) were injected into the 4^th^ mammary fat pad of 7-8 weeks old female BALB/c mice. Treatments were initiated 7 days after tumor inoculation. A depilatory cream was used to remove the fur at the tumor site to enable accurate measurement of the tumor dimensions. Mouse survival, body weight changes, and tumor growth were monitored and recorded every other day (eod). The tumors were measured using vernier calipers, and the tumor volume was calculated as V = ½ x L x W^2^. The tumor growth inhibition (TGI) was calculated as follows: TGI = (1 - mean final tumor volume of treatment group/mean final tumor volume of the control group)*100. Animals exhibiting signs of distress such as hunched posture, ruffled fur, reduced mobility, and greater than 15% weight loss were euthanized via carbon dioxide intoxication.

### Treatment regimens

#### Varying TGFβi dosing frequencies

Mice were randomized into groups of 4 and were treated as follows: (i) normal saline intravenously (i.v.) four times over 2 weeks (Th, M, Th, M) and intraperitoneally (i.p.) daily (Th-M) for 2 weeks; (ii) 32 mg/kg SB-POx i.p. daily (Th-M) for 2 weeks; (iii) 32 mg/kg SB-POx i.p. eod (Th, Sat, M) for 2 weeks; (iv) 75 mg/kg PTX-POx i.v. four times over two weeks (Th, M, Th, M); (v) combination of ii and iv; (vi) combination of iii and iv.

#### Varying TGFβi dose and comparison of i.v. versus i.p

Mice were randomized into groups of 4 and were treated as follows: (i) normal saline i.v. four times over 2 weeks (Th, M, Th, M) and i.p. eod (Th, Sat, M) for 2 weeks; (ii) 50 mg/kg SB-POx i.p. eod (Th, Sat, M) for 2 weeks; (iii) 50 mg/kg LY-POx i.p. eod (Th, Sat, M) for 2 weeks; (iv) 75 mg/kg PTX-POx i.v. four times over two weeks (Th, M, Th, M); (v) Combination of ii and iv; (vi) Combination of iii and iv; (vii) SB/PTX-POx (5.2/8) i.v. four times over 2 weeks (Th, M, Th, M); (viii) LY/PTX-POx (5.2/8) i.v. four times over 2 weeks (Th, M, Th, M); (ix) LY/PTX-POx (4/8) i.v. four times over 2 weeks (Th, M, Th, M).

#### Oral dosing of TGFβi

Mice were randomized into groups of 5 and were treated as follows: (i) normal saline i.v. four times over 2 weeks (Th, M, Th, M) and via oral gavage (o.g.) eod (Th, Sat, M) for 2 weeks; (ii) 50 mg/kg SB-POx o.g. eod (Th, Sat, M) for 2 weeks; (iii) 50 mg/kg LY-POx via o.g. eod (Th, Sat, M) for 2 weeks); (iv) 75 mg/kg PTX-POx (injected i.v. once every 4 days for two weeks); (v) Combination of ii and iv; (vi) Combination of iii and iv. In addition to the POx formulations, SB and LY were also administered in the form of a suspension composed of 0.5% sodium carboxymethylcellulose (NaCMC) and 0.25% Tween 80 as the control formulation.

### Micro-computed tomography (micro-CT)

Animals from three groups (Saline (i.p. & i.v.), SB-POx 50mg/kg (i.p.), and SB-POx 50mg/kg (i.p.) & PTX-POx 75 mg/kg (i.v.)) were chosen for whole-body micro-CT. The animals were anesthetized in an inhalation induction chamber (3% isoflurane in oxygen) and placed in the micro-CT chamber in the supine position (anesthesia maintained at 2.5% isoflurane). The images were acquired on a PerkinElmer Quantum-GX2 Imaging System (PerkinElmer, Inc., Waltham, MA USA) with a 120 µm voxel size using the following scan settings: x-ray tube voltage of 90 kV, current of 88 µA, Cu[0.06mm]+Al[0.5mm] filter, 4 min scan duration, and a 60 mm FOV. The scans were subjected to the retrospective lung-gating protocol, creating two datasets corresponding to the inspiration and expiration phases of the breathing cycle. All datasets were calibrated to the Hounsfield scale. The image sets from the inspiration phase was used for analysis. A qualitative inspection was performed for each dataset to determine the presence of any metastatic lesions or tumors. The total lung volume was then segmented for each animal using a uniform threshold of - 400 HU to capture the lung tissue while excluding the vasculature. The mean lung tissue density (HU) over the entire volume was measured and compared across all study groups. Differences between groups were used to further determine the presence of potential lesions. All analyses were performed using Analyze 14.0 software (AnalyzeDirect, Inc., Overland Park, KS USA).

### Histology

The resected lungs were fixed in 10% formaldehyde buffered solution for ≥ 24h, followed by routine serial dehydration and paraffin embedding. The lungs were sectioned to 5 µm, stained with routine hematoxylin-eosin, and scanned using the Aperio AT2 digital scanner (2021 Leica Biosystems Imaging, Inc., Deer Park, IL, USA) at 20X magnification. The micrometastases in the lung were quantified using an image analysis algorithm. A boarded veterinary pathologist working in the UNC PSC used Definiens Architect XD 64 version 2.7.0.60765 to identify tumors in the sections, including those within and adjacent to lung parenchyma (presumed tumors arising from the thoracic lymph nodes). First, tissue (ROIs) on each slide was automatically detected using 221 brightness and 3.6 homogeneity thresholds and a minimum tissue size of 150,000 um^2^. The two tissue sections on the slide were randomly assigned ROI labels “tissue 1” or “tissue 2” by the computer software. Next, large vessels, bronchi, and thymus were manually excluded from the ROIs. Examples of lung, glass (alveolar spaces), blood (erythrocytes in large vessels or alveolar spaces), and tumor were provided to the analysis software, which categorized the remaining lung sections into the appropriate tissue category. The resulting tumor area (um^2^) was then divided by the lung area (um^2^) for each ROI. Values were presented as tumor percent of the lung. The algorithm was validated by a randomized, semi-quantitative assessment of the relative percent tumor burden by the same veterinary pathologist blinded to treatment groups.

### Genomic Data Processing and Visualization

Upper quartile normalized mRNA sequencing data from Hollern et al. [19](GEO: GSE124821) and from Unpublished et al. (GEO: GSE223630) were merged to a combined datatset of 232 individual mouse mammary tumors and normal mammary gland controls. Tumor models that contained less than three biological replicates were filtered out prior to log2 transformation and median centering by gene. Genes of interest were plotted by tumor model using the ggplot2 package (Version 3.3.6).

### Data availability

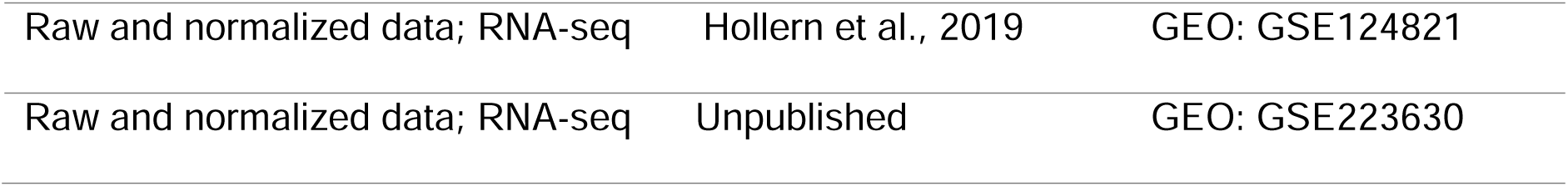

## Results

### Characterization of SB and LY containing PMs

The POx PMs loaded with SB at 8/20 drug/polymer mass ratio were small with a reasonably narrow size distribution as evident from DLS (34 nm, PDI 0.17) and had a somewhat asymmetrical but compact shape based on TEM (**Fig. 1**). The PTX-containing PMs were also small and even more uniform (28 nm; PDI 0.07) which is consistent with our previous publication [20]. Co-loading of these two drugs in PMs at 5.2/8/20 SB/PTX/POx mass ratio produced uniform particles (37 nm; PDI 0.07) as well. On the other hand, PMs loaded with LY had a bimodal size distribution, with a smaller peak at around 18 nm and a larger peak at 110 nm (**Fig. 1**). The mixture of small and large particles was also visible in TEM images. The co-loaded LY/PTX-POx were prepared at two mass ratios LY/PTX/Pox 5.2/8/20 and 4/8/20. Of the two formulations, one (4/8/20) formed small micelles (30.6 nm; PDI 0.18) that had a nearly spherical shape (**Fig. 1**). The other (5.2/8/20) formed relatively large particles (∼140 nm; PDI 0.19) that appeared to be a mixture of separate and flocculated spheres, some of which may have further coalesced into the worm-like structures (**Fig. 1**).

**Fig. 1.**
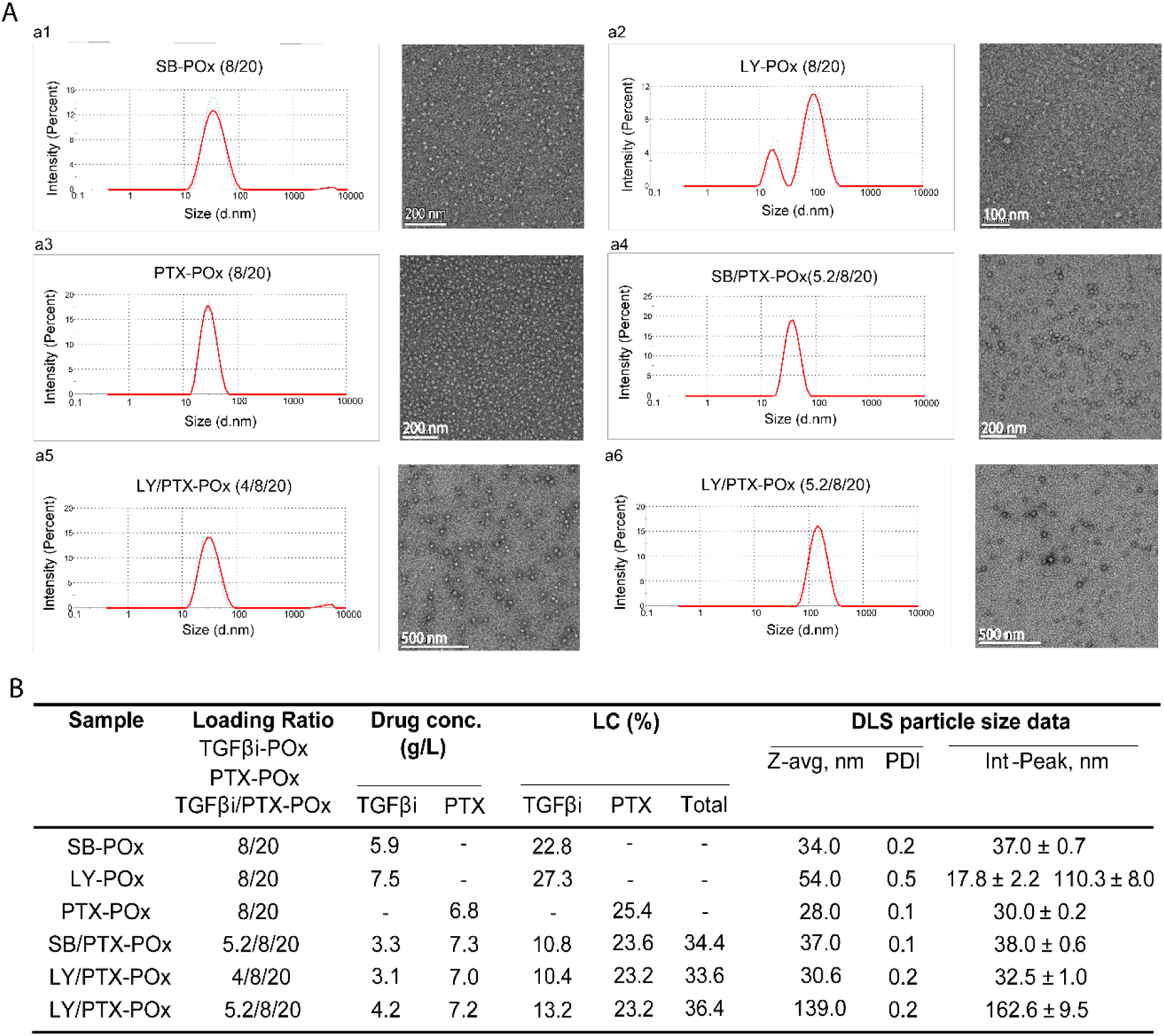
Characterization of single TGFβi and co-loaded TGFβi/PTX in POx micelles by DLS and TEM. **(A)** DLS intensity size distributions and the corresponding TEM images of the nanoassemblies formed in drug-loaded PMs of various compositions (a1) SB-POx (8/20); (a2) LY-POx (8/20); (a3) PTX-POx (8/20); (a4) SB/PTX-POx (5.2/8/20); (a5) LY/PTX-POx (4/8/20); (a6) LY/PTX-POx (5.2/8/20). The numbers in the brackets represent the mass ratios TGFβi/POx, PTX/POx or TGFβi/PTX/POx. **(B)** Drug concentration, loading capacity (LC (%) = M_drug_ / (M_drug_ + M_excipient_) × 100 (%)) and DLS characteristics (z-average hydrodynamic diameter, polydispersity index (PDI), diameters at intensity size distribution maxima) of the formulations presented in (A). Drug concentration in the solution was measured by HPLC. For DLS measurements, the samples prepared at POx 20 mg/ml were diluted 10 times.

### TGF**β** inhibition activity of SB and LY and *in vitro* cytotoxicity of SB, LY, and PTX

TGFβ modulates ECM remodeling of the tumor microenvironment (TME) by activating TGFβ signaling in stromal cells, particularly fibroblasts, which engage in pro-fibrotic responses [21]. Phosphorylation of SMAD2 and SMAD3 are hallmarks of TGFβ signaling (**Fig. 2A**). We used p-SMAD2/3 expression levels as a marker for assessment of the inhibitory effect of our drugs on TGFβ signaling in TGFβ-stimulated NIH-3T3 fibroblasts. The western blot and flow cytometry analysis revealed p-SMAD2/3 attenuation in groups treated with TGFβi relative to control groups. The TGFβ inhibition appeared to be more pronounced in groups treated with LY than SB, although the western blot quantification presented a trend but no significant difference between these groups (**Fig. 2B, 2C**). However, the flow cytometry also corroborates the conclusion that LY is more potent than SB (**Fig. 2D, 2E**). This difference may be explained by the ability of LY to inhibit both TGFR-I&II, whereas SB inhibits only TGFR-I [22]. Both vehicles, DMSO and POx, had a negligible effect on the p-SMAD2/3 levels. Because of the dose-dependent cytotoxicity of PTX (IC50 0.7 nM, Supplementary **Fig. S1**), we did not include this drug in the TGFβ inhibition study. Notably, SB and LY were substantially less toxic than PTX at concentrations ranging from 0.256 nM to 100 μM (Supplementary **Fig. S1**).

**Fig. 2.**
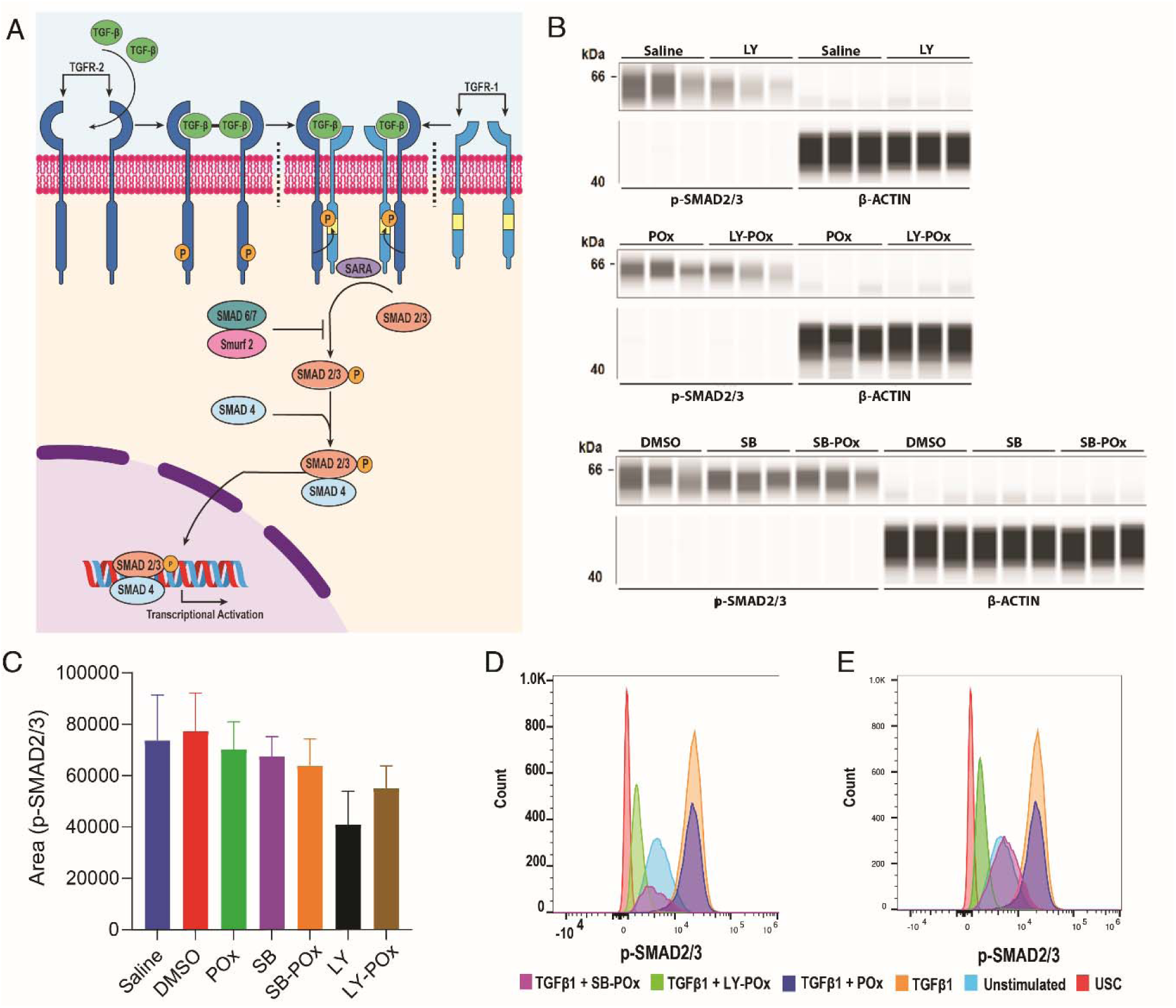
TGFβi treatment suppresses TGFβ signaling in NIH-3T3 fibroblasts. (**A**) Schematic of canonical TGFβ signaling. (**B**) Capillary-based western blot analysis of expression of p-SMAD2/3 (∼62kDa) and housekeeping protein, β-ACTIN (∼48kDa) in NIH-3T3 cells treated with TGFβi in POx PMs (SB-POx (8/20) and LY-POx (8/20)) or dissolved in DMSO. Control groups were treated with the same amounts of POx or DMSO. The images were generated using Compass for Simple Western (version 6.0). (**C**) The quantification of p-SMAD2/3 bands in **B**. *p<0.05 computed by one-way ANOVA with Tukey’s multiple comparisons test. (**D, E**) The p-SMAD2/3 expression analyzed by flow cytometry in NIH-3T3 cells treated with TGFβi **(D)** formulated in POx PMs or **(E)** dissolved in DMSO. The D and E panels were run in the same experiment and split for the clarity of presentation to avoid overlap between the free and micelle-formulated drug histograms. The panels have common histograms for unstained (USC), unstimulated, TGFβ stimulated controls.

### Evaluation of TGF**β**i and their combinations with PTX in the 4T1 immune-competent animal model of TNBC

It has been previously reported that TGFβ downregulation by TGFβi is sustained for only a few hours after the drug is cleared from circulation, suggesting that frequent injection of LY is necessary for the persistent TGFβ pathway suppression [23]. Accordingly, we decided to test daily versus every-other-day (eod) i.p. dosing regimens for one formulation of TGFβi (SB-POx). To this end, we used the 4T1 immune-competent mouse model of TNBC, which is highly metastatic, therefore allowing for the testing of the treatment effectively both on primary as well as metastatic tumor burden. The tumor inhibition effect of SB-POx at 32 mg/kg with respect to the primary tumor was not significantly different between the two regimens (**Fig. 3A).** Of note, i.v. administration of PTX-POx alone at 75 mg/kg suppressed the primary tumor growth relative to the control (**Fig. 3A** and **3B**). The tumor inhibition of the combination of SB-POx (i.p.) and PTX-POx (i.v.) did not differ significantly from that of PTX-POx (**Fig. 3A** and **B**). As far as the lung macro- and micrometastases were concerned, the differences between any of these groups were not significant. However, there was a trend for a decrease in the metastatic spread for PTX-POx treated mice and even more so for SB-POx using the eod regimen (**Fig. 3C** and **3D**). The animals maintained their weight during the course of the treatments (**Fig. 3E**).

**Fig. 3.**
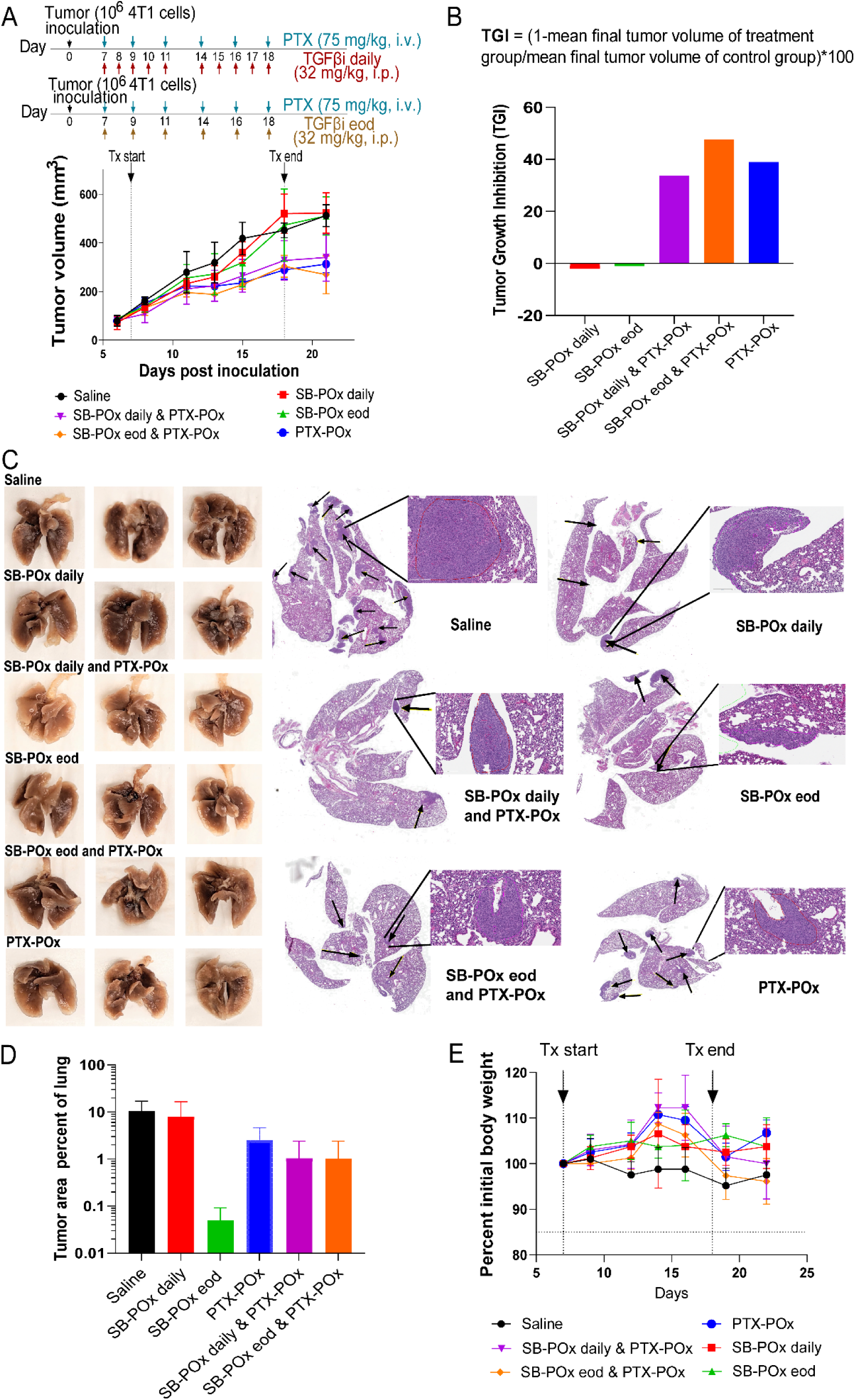
TGFβ inhibition synergizes with chemotherapy to inhibit primary tumor growth and lung metastases in 4T1 TNBC tumor-bearing mice. (**A**) Tumor growth curves following treatments with single drug SB-POx (8/20) i.p., single drug PTX-POx (8/20) i.v., or separately administered combinations SB-POx (8/20) i.p. and PTX-POx (8/20) i.v. using different SB-POx schedules (daily or eod) at 32 mg/kg. See supplementary Table S3 for the complete statistical comparison between all groups. (**B**) Percent tumor growth inhibition corresponding to tumor growth curves. (**C**) Images of lungs with metastatic nodules (left) and corresponding H&E stain images (right) with metastatic nodules indicated with black arrows from representative mice in each group. (**D**) Histopathological scoring of lung metastatic burden. (**E**) Body weight changes (percent of initial) in mice treated with TGFβi-POx and/or PTX-POx.

After determining that the eod dosing regimen for TGFβi was as effective as daily dosing, we decided to adhere to it and increased the TGFβi dose to 50 mg/kg for both SB and LY. This improved the performance of SB-POx alone, which slowed down tumor growth relative to the saline control (**Fig. 4A**). LY-POx also slowed tumor growth but to a lesser extent than SB-POx. Consistent with the previous experiment, i.v. PTX-POx produced a notable anti-tumor effect. Moreover, we observed that the combinations of PTX-POx with either TGFβi formulated in PMs provided some benefit over PTX-POx alone (**Fig. 4C**). There was a trend for an increase in anti-tumor effect when the PTX-POx was combined with either LY-POx or SB-POx. Because the metastatic burden of the saline-treated controls was at the low end, possibly due to the poor metastasis formation in the two animals selected for histopathology from this group, the effect of the treatments on the metastatic burden was confounded (**Fig. 4D** and Supplementary **Fig. S3**). Nonetheless, compared to the saline control in **Fig. 3D**, the single drug treatments, as well as drug combinations, appeared to lower the metastatic burden in the lungs.

**Fig. 4.**
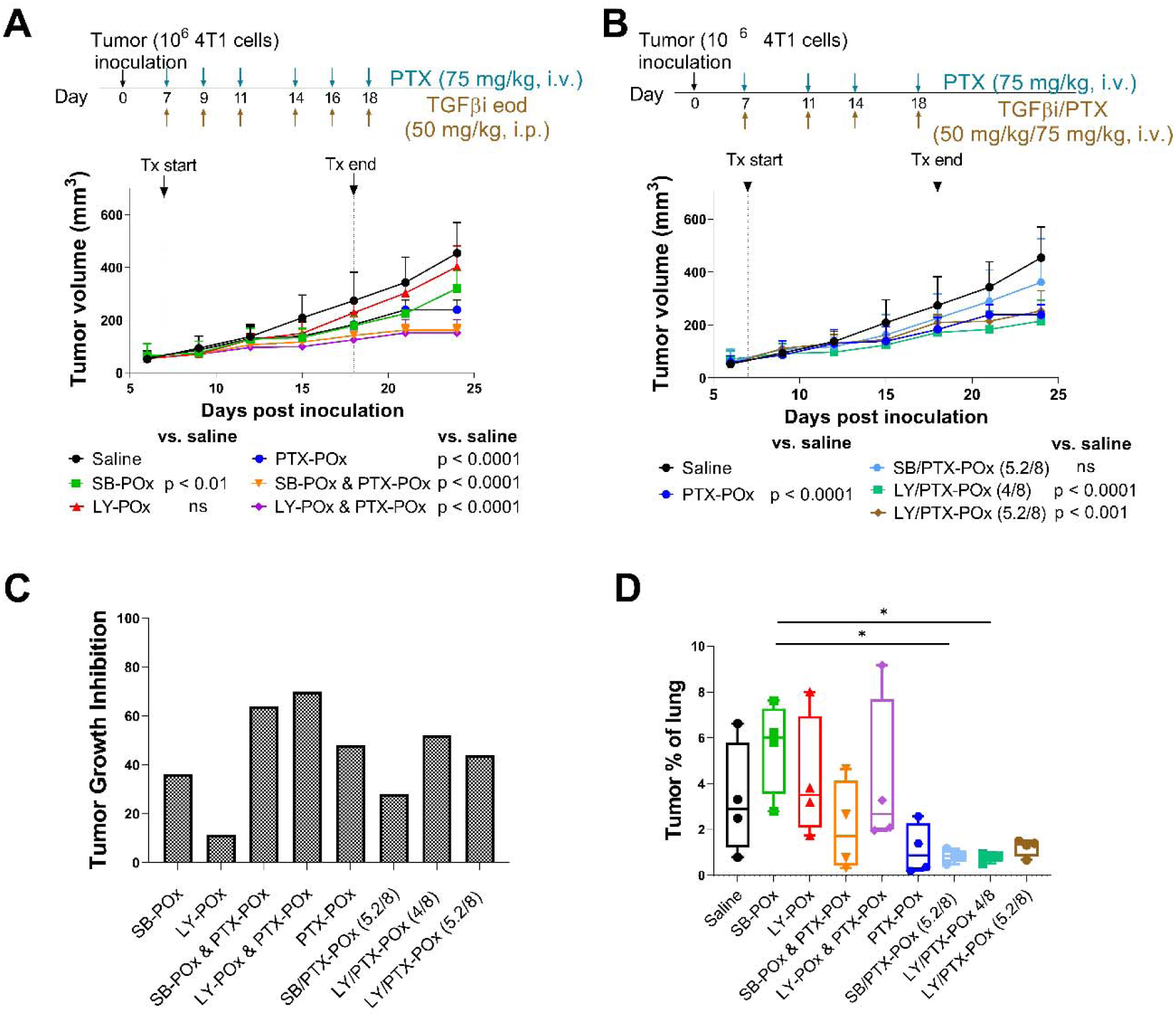
Tumor growth inhibition with i.p. versus i.v. routes of TGFβi delivery. (**A**) Tumor growth curves in 4T1 tumor-bearing mice treated with either single drug or separately administered TGFβi and PTX combinations, or (**B**) co-loaded micelles of TGFβi and PTX (common saline and PTX-POx groups were used for experiments 4A and 4B since the experiments were conducted in parallel); See supplementary Table S4 for the complete statistical comparison between all groups. (**C**) Percent tumor growth inhibition by treatments corresponding to 4A and 4B. (**D**) Histopathological scoring of lung metastatic burden. Data represent mean ± SD. N=2 with duplicate sections. *p < 0.05 computed by one-way ANOVA with Tukey’s posthoc test.

To reaffirm the extent of metastatic spread in the lungs of live animals in these groups, we examined the lungs of the tumor-bearing animals for metastatic lesions by the *in vivo* micro-CT. The lesions were small and below the resolution limit of this method (Supplementary **Fig**. **S2A**). Nonetheless, the overall lung tissue density appeared to decrease between saline, single SB-POx, and combination SP-POx and PTX-POx groups as estimated by the Hounsfield units (HU) (HU −594, −606, −636, respectively) (Supplementary **Fig. S2B**). Although these differences were minor and not significant, they suggest a trend for decreasing metastatic nodules. The animal body weight was maintained during the single drug treatments. It decreased by ∼13% during the combination drug treatments, although even in this case, it still stayed within the 15% cut-off limit by the end of the study (Supplementary **Fig. S2C**).

Since the co-loading of several drugs in PMs previously resulted in considerable treatment benefits [24], we examined the co-loading approach for the combination of TGFβi and PTX. Here, we wanted to see if a co-delivery of these agents in the same nanocarrier provided any benefit over separate administration of TGFβi and PTX at the same drug dose. The two examined co-loaded drug formulations, LY/PTX-POx (5.2/8/20) and LY/PTX-POx (4/8/20), administered i.v. displayed significant anti-tumor activity, while one, SB/PTX-POx (5.2/8/20), was statistically not different from the saline control (**Fig. 4B**). The tumor growth inhibition in these treatment groups did not exceed ca. 50% even with the most potent combinations (**Fig. 4C**). None of these combinations surpassed the efficacy of single PTX-POx, and none of them was as effective as the combination treatments with micellar TGFβi and PTX administered separately (**Fig. 4A** and **Fig. 4C**). This could be the effect of the cumulative dose of TGFβi which was 200 mg/kg for co-loaded drugs (four injections, i.v.) and 300 mg/kg for drugs administered separately (six injections, i.p.). Since frequent i.v. injections cannot be realized in mice due to the risk of damaging the tail vein with repeated needle punctures, the eod regimen for the co-loaded micelles was not tested for the i.v. route. We also would like to point out that the co-loading strategy appeared to have some effect against the metastatic spread in the lungs, as evidenced by the lower percentages of metastatic burden in these groups. However, none of the values were significantly different from the saline control (**Fig. 4D** and supplementary **Fig. S3**).

Finally, we tested the oral route of delivery for TGFβi using the eod regimen because both SB and LY are orally available [25, 26]. In addition to POx formulations, SB and LY were also formulated in a conventional vehicle for oral administration (0.5% NaCMC and 0.25% Tween 80) to evaluate if POx micelles offer any advantage in terms of oral drug delivery. The results are presented in **Fig. 5**. The single TGFβi was not effective when administered via o.g., independent of the formulation. In one case, o.g. SB-POx, significantly increased tumor growth. (That contrasted sharply with the i.p. SB-POx administered at the same dose (50 mg/kg, **Fig. 5A and 5B**), which produced notable tumor inhibition.) The combination treatments with o.g. TGFβi and i.v. PTX-POx, in all but one case, produced significant inhibition of the tumor growth compared to saline controls. However, they were not statistically different from the single i.v. PTX-POx treatments. In this study, the o.g. TGFβi and i.v. PTX-POx combination, as well as other treatments, produced a significant decrease in lung metastasis, as evidenced by the dramatic decrease in the metastatic lesions in the lungs of mice treated with the drugs when compared to the saline-treated mice (**Fig. 5C**). Overall, when administered orally in combination treatments, the TGFβi in PMs performed as well as these agents in the conventional vehicle. Except for minor fluctuations, the animals in all the groups maintained their weight (Supplementary **Fig. 5D**). However, we need to point out that the animals receiving combination treatments with the oral TGFβi in the conventional vehicle were visually less active and appeared more dehydrated (displayed ruffled fur) than the groups receiving combinations with oral TGFβi in PMs.

**Fig. 5.**
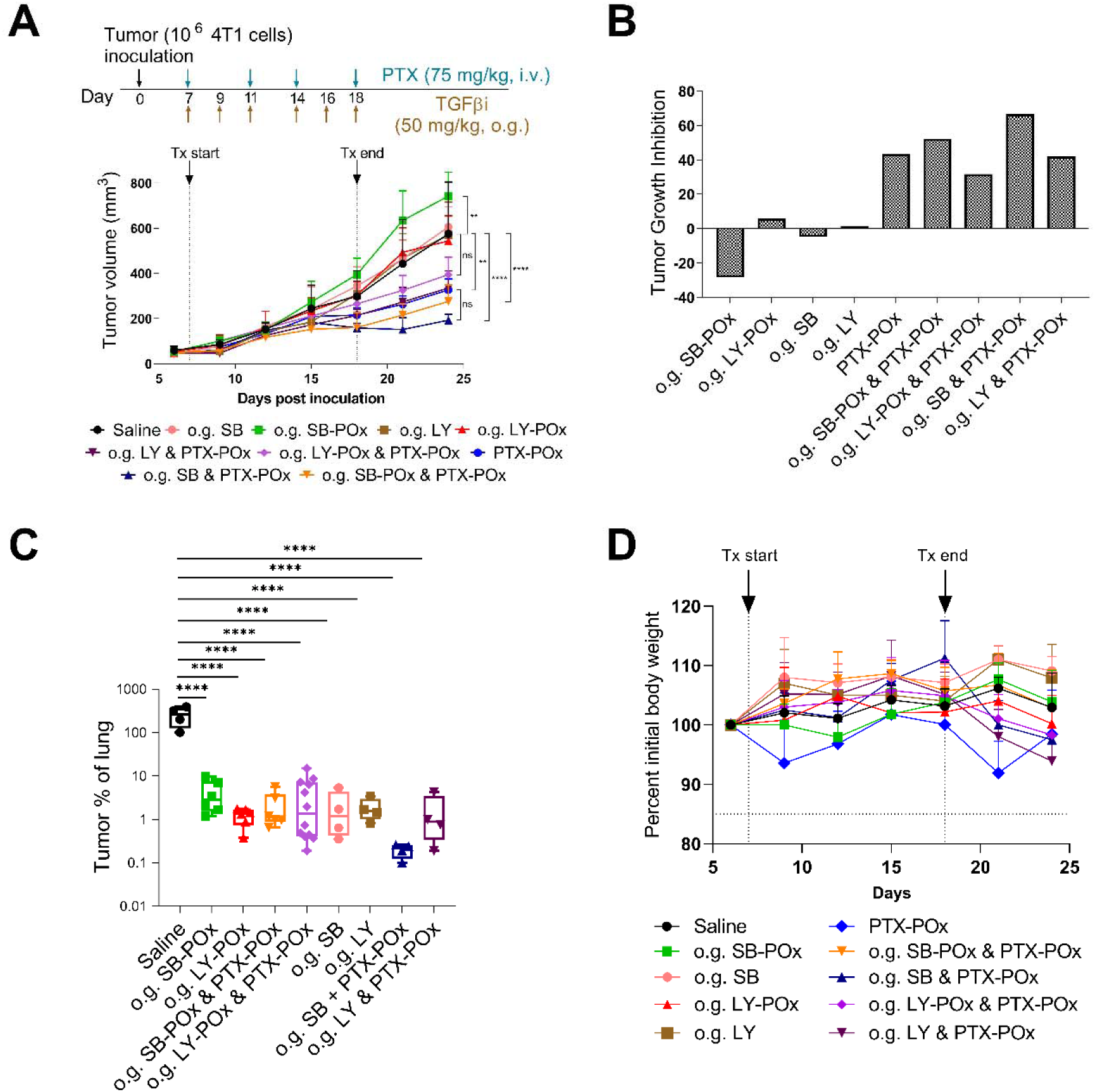
Tumor growth inhibition with oral route of TGFβi delivery. (**A**) Tumor growth curves in 4T1 tumor-bearing mice treated with orally delivered (o.g.) TGFβi formulated either in POx or 0.5% NaCMC and 0.25% Tween 80. Data represent mean ± SD. N=4. **p < 0.003, ****p < 0.0001 computed by two-way ANOVA with Tukey’s posthoc test. See supplementary Table S5 for the complete statistical comparison between all groups. (**B**) Percent tumor growth inhibition by treatments corresponding to 5A. (**C**) Histopathological scoring of lung metastatic burden. Data represent mean ± SD. N=2 with duplicate sections. ****p < 0.0001 computed by one-way ANOVA with Tukey’s posthoc test. (**D**) Body weight changes (percent of initial) in mice treated with TGFβi-POx and/or PTX-POx.

To further explore the possibility of using the TGFβi and PTX PMs to treat other TNBC models, we screened an array of gene expression profiles of 19 transplantable syngeneic genetically engineered mouse (GEM) models of TNBC and two normal mammary gland controls available at UNC Lineberger Cancer Center and assessed them based on the relative expression of genes known to regulate both the canonical and non-canonical TGFβ pathways (*Tgfbr1, Tgfbr2, Tgfb1, Tgfb2, Vim, Cd36, Egfr, Mapk1, Mapk3, Pik3ca*). As presented in Supplementary **Fig. S5**, our 4T1 model displays TGFβ signature genes (*Tgfbr1, Tgfb1*), EMT signature (*Vimentin and Mapk3*), and fibrotic signature (*Pik3ca*), and a low expression of fibroblast quiescence marker (*Cd36*). We selected two additional tumor models, T11-UV and T11-Apobec, that displayed a high expression of *Tgfb1* (both tumor models) and *Tgfb2* (T11-UV), high expression of *Vimentin* (*Vim*) and fibrotic marker (*Pik3ca*), and low expression of *Cd36* and apoptosis regulator *Bcl2* (Supplementary **Fig. S5**). Both models are derived from T11 murine breast cancer that closely resembles the genomic phenotype and immune responses seen in patients with Claudin-low breast cancer [19, 27–29]. T11 cells (a *Trp53*-/-cell line) overexpressing *Apobec3* and T11 cells exposed to short-wave UV exposure were transplanted into syngeneic hosts to generate T11-Apobec and T11-UV models, respectively [19]. Interestingly, these models responded differently to our drug treatments. TGFβi SB-POx or LY-POx alone markedly inhibited the T11-Apobec tumor growth (**Fig. 6A, C**). The PTX-POx alone had a significant but smaller tumor inhibition effect. The tumor growth regressed more significantly when treated with the combination of TGFβi-POx and PTX-POx compared to either of the treatments alone. In contrast, in the T11-UV tumor, the response to the single drug TGFβi therapies SB-POx or LY-POx was considerably less than that observed in single drug PTX-POx treatment (**Fig. 6B, D**). In this regard, T11-UV behaved somewhat similarly to the 4T1 tumor. The combinations of TGFβi and PTX PMs did not provide an added benefit in these tumors over the PTX-POx treatment. All animals maintained their weights during the treatments (**Fig. 6E, F**).

**Fig. 6.**
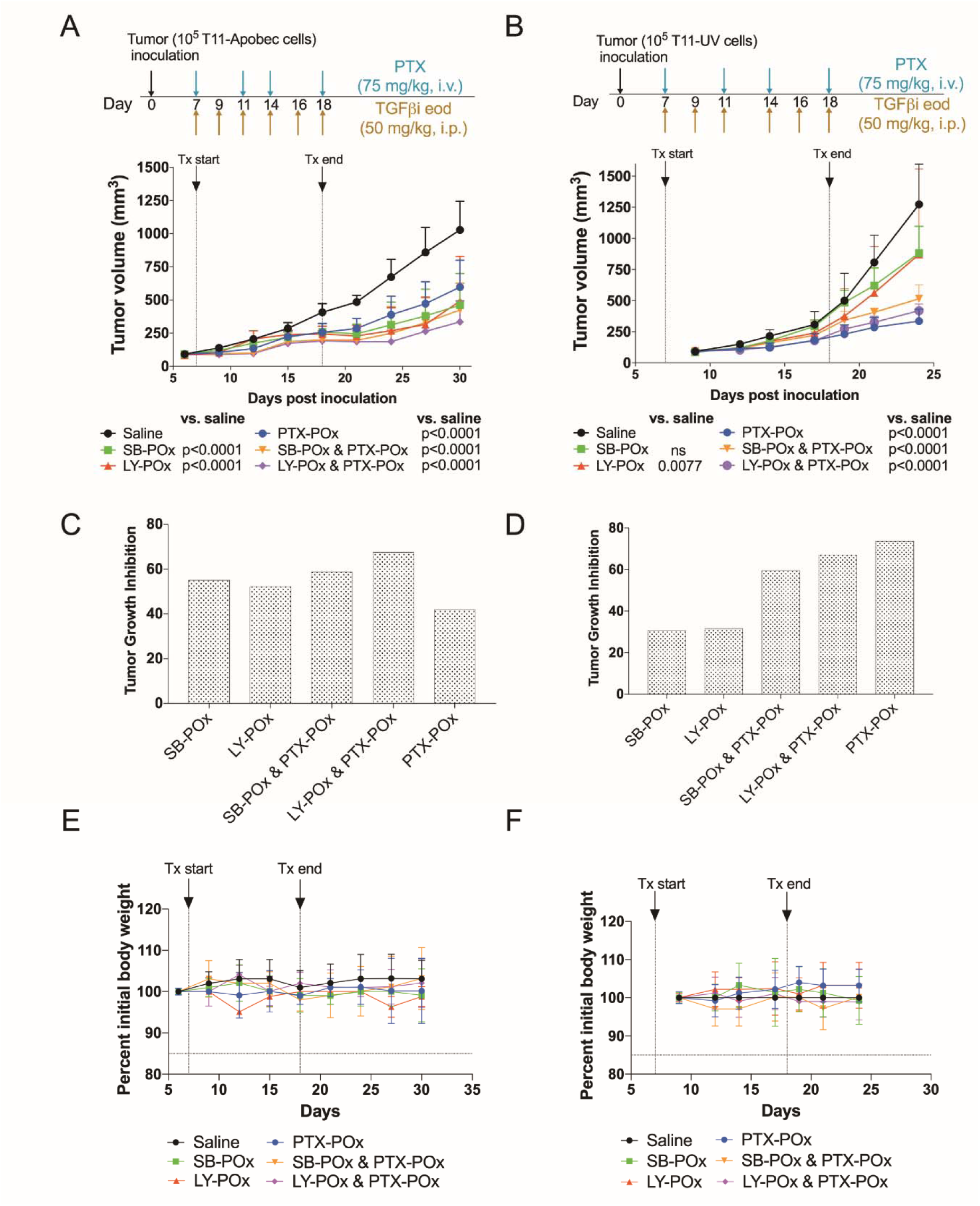
Differential sensitivity of Claudin-low TNBC subtypes to TGFβi and PTX. Tumor growth curves following treatments with single drug SB-POx (8/20) i.p., single drug PTX-POx (8/20) i.v., or separately administered combinations SB-POx (8/20) i.p. and PTX-POx (8/20) i.v. in mice bearing **(A)** T11-Apobec and **(B)** T-11 UV tumors. Data represent mean ± SD. N=4. **p < 0.003, ****p < 0.0001 computed by two-way ANOVA with Tukey’s posthoc test. See supplementary **Tables S6** and **S7** for the complete statistical comparison between all groups. (**C, D**) Percent tumor growth inhibition by treatments corresponding to 4A and 4B as determined at the endpoint of tumor growth experiments. (**E**, **F**) Body weight changes (percent of initial) in mice treated with TGFβi-POx and/or PTX-POx.

## Discussion

POx micelle nanoformulation technology demonstrates an extremely high loading of small molecules while requiring a significantly lesser amount of polymeric excipient than conventional vehicles [30]. This translates to 1) high-dose PTX delivery with few side effects [20] and 2) co-delivery of two drugs in a single micelle, thereby increasing the tumor distribution and anti-tumor effect of these drugs [24, 31]. Our goal in this study was to evaluate whether this platform technology could improve treatment outcomes during chemo-immunotherapy in the mouse models of TNBC. Here we combine PTX, one of the most active cytotoxic agents for TNBC medical management [32], with the agents capable of inhibiting the TGFβ signaling pathway.

The TGFβ cytokine family comprises three isoforms, TGFβ1, TGFβ2 and TGFβ3, two transmembrane serine/threonine receptors, TGFR-I, TGFR-II and a coreceptor, TGFR-III. The cytokines are homodimers that initiate the TGFβ signaling by binding to TGFR-II, which is constitutively active and undergoes autophosphorylation. The phosphorylated TGFR-II recruits TGFR-I resulting in the formation of a heterotetrameric complex of TGFR-I and TGFR-II, following which TGFR-II phosphorylates the serine residues of TGFR-I [33]. The signal is relayed downstream through the phosphorylation of receptor-regulated SMADs (R-SMADs: SMAD 1, SMAD2, SMAD 3, SMAD 5 and SMAD 9) by TGFR-I; subsequently, the phosphorylated R-SMADs form a complex with co-SMAD and the complex translocates to the nucleus and initiates gene transcription [34].

Clinical studies of small molecule TGFβi involve their repeated oral administration, either twice or once daily, due to the short-lived effect of p-SMAD inhibition by these molecules [35]. In the present study, we varied dosing routes and frequencies to test if i.p. administration of TGFβi improved its anti-tumor efficacy compared to o.g. The i.v. administration route is not suitable for repeated dosing of drugs and was not included in our design since it is closely approximated by the i.p. administration in rodents [36].

Two small-molecule TGFβi with different potencies for the kinase domain were studied in this work: 1) SB (GlaxoSmithKline), an imidazole derivative with an IC_50_ of 14.3 nM for TGFR-I; and 2) LY (Eli Lilly), a pyrazole-based dual inhibitor of TGFR-I (Ki = 38 nM) and TGFR-II (Ki = 300 nM) [22]. Both agents were tested in animal models (rodents), but neither advanced to human clinical evaluation, likely due to their poor solubility. To our knowledge, none of them has a format available for either i.v. or p.o. administration. We successfully reformulated these molecules in our POx PMs, resulting in highly loaded, small, and stable nanoparticles for SB and somewhat less stable but still useful nanoformat for LY. The resulting POx nanoformulations in the present study generally demonstrated safety and activity, which provides a basis for further translational development of these agents.

We used a well-known 4T1 TNBC model in the initial phase of our work. According to a recent study by our group, the 4T1 model displays an intermediate gene expression pattern that is more like a hybrid between these two subtype states, lacking clear Claudin-low or basal-like transcriptomic features [37], which contradicts previous literature that suggests this model is simply basal-like [38–40]. The 4T1 tumors display an immune suppressive gene signature with upregulation of Th2, T-regulatory, PD-1, and CTLA4 signaling [41]. As for the mutation signature, a multi-omics study of 4T1 tumors reported mutation in the Trp53 and PIK3cg (catalytic subunit of PI3 kinase) genes. The study also revealed mutations in markers for metastasis, cell proliferation, and cancer-related genes. The 4T1 model is highly aggressive and does not display a mutation in the BRCA genes [41]. BRCA-deficient basal-like tumors are more sensitive to chemotherapy. Because TGFβ signaling promotes the development of T-regulatory cells, which are key players in regulating immune tolerance, it strengthens the rationale for using TGFβi in this tumor to restore the tumor’s suppressed immune surveillance [42].

Since the i.p. route has a greater extent and rate of absorption than the oral route, we examined if daily dosing of TGFβi in PMs has any benefit compared to the eod dosing [36]. The tumor inhibition study using the 4T1 TNBC model revealed that daily dosing of micellar SB-POx was comparable to eod dosing with regard to the primary as well as metastatic tumor inhibition. Based on this, we proceeded with the eod dosing. Of the two agents, SB-POx appeared to be more effective – at the higher dose, it has shown potency in inhibiting primary tumor growth. In contrast, LY-POx had a modest effect on the primary tumor. Regarding suppression of lung metastases, a secondary endpoint in our study, we did not see definitive evidence of the superiority of any of our formulations. However, we observed at least a trend of improved suppression with several treatments, which provides the basis for further assessment of this matter using dedicated metastatic models. The effect of the TGFβi on the metastatic spread could be related to the suppression of the TGFβ-mediated EMT program, which can reduce the invasive and metastatic capacity of the primary tumors [43].

We further evaluated if the oral dosing of TGFβi in either a conventional vehicle or micelles could be more advantageous than systemic POx formulation. As a single drug, there was no benefit of oral dosing, and for one drug, SB, the oral dosing was even detrimental and increased tumor growth. In contrast, SB-POx (i.p.) at the same dose produced significant tumor inhibition. The underperformance of the oral SB-POx compared to the systemic dosing could be due to the poor oral absorption of the drug, resulting in its sub-therapeutic levels in the tumor. The interspecies difference in the oral bioavailability of TGFβi has been previously reported, with considerably lower bioavailability in small animals compared to large animals [44]. Furthermore, prolonged exposure to a low dose of TGFβi has been shown to result in the development of drug resistance [23]. This could be attributed to the glycoprotein P (P-gp) expression in the intestinal endothelium in the gut, which could be overcome by P-gp inhibitors such as Tween 80 contained in the conventional vehicle NaCMC [45]. However, Tween 80 is implicated in several side effects and, therefore, must be substituted by alternative, better-tolerated formulations [46].

Next, we proceeded with the combination therapy using TGFβi and PTX. First, we concluded that the co-formulation of these agents in the same polymeric micelle does not have any benefit compared to their separate administration because of the very different pharmacodynamics and treatment schedules of these drugs. The drugs administered separately at their optimal dosing schedules have shown superior tumor inhibition compared to co-formulated drugs. Second, we clearly see that the combination treatments with two drugs surpass treatments with each single TGFβi in the 4T1 model. The combination treatments using the higher dose of TGFβi also appear to be more effective than the single PTX-POx. For the combination treatments within the design of this study, we could not see any significant difference between the oral and systemic dosing of TGFβi PMs.

To further evaluate the broad applicability of our dosing approach, we tested the anti-tumor efficacy of TGFβi and PTX in POx PMs using our credentialed mouse models of TNBC. We selected two Claudin-low TNBC models, T11-Apobec and T11-UV, based on the expression of genetic markers for TGFβ signaling and EMT. Most importantly, the combination treatments using these two drugs in our PMs format were uniformly effective across all three models, including the 4T1 model discussed above. However, we noted with interest that the T11-Apobec and T11-UV models showed different sensitivities to single-drug treatments. Specifically, the T11-UV tumor responded well to PTX-POx but had lower responses to either of the TGFβi-POx. Conversely, the T11-Apobec tumor was highly responsive to each of the single TGFβi-POx but was somewhat less responsive to PTX-POx.

We would like to point out the differences in the mutation signatures and tumor mutation burden (TMB) of the T11-Apobec and T11-UV tumors, which are known to regulate response to treatments. The mutation signatures of cancers dictate their immunogenicity and, thus, response to treatments that stimulate immune surveillance, such as immune checkpoint blockade [47, 48]. Apobec3 is a family of antiviral enzymes that are implicated in tumorigenesis. Tumors overexpressing *Apobec3* are less likely to respond to chemotherapy but are sensitized to immunotherapy [49]. Moreover, *Apobec3G* expression correlates with the expression of TGFR1 and TGFβ1, one (TGFβ1) of which was found to be highly expressed in this model. In addition to Apobec3G regulating TGFβ1 signaling [50], the latter also regulates the former’s expression [51], alluding to possible crosstalk between the pathways. Therefore, targeting the TGFβ pathway should also attenuate Apobec3-mediated resistance of tumors to chemotherapy. The observed synergistic effect of the combination of TGFβ and PTX in these tumors in the present study further corroborates this.

The transcriptome evaluation of the Claudin-low T11-UV tumors revealed a decrease in the *Bcl2* gene expression compared to the normal mammary tissue. *Bcl2* is an apoptosis regulator in cells, and its expression negatively correlates with sensitivity to PTX [52]. We noted that the gene expression of *Bcl2* in T11-UV was even lower than in T11-Apobec, which could explain the higher sensitivity of this tumor to PTX relative to T11-Apobec. Also, T11-UV has a higher mutation load than T11-Apobec (TMB = 1783 and 374, respectively, and both are markedly higher than the parental cell line, T11 (TMB = 117) [19]). Higher TMB for many human tumors has been shown to correlate with higher levels of neoantigens and with increased survival after immune checkpoint inhibitor therapy [53, 54]. Since TGFβ controls the regulation of immune tolerance [55] and response to immunotherapy [56], higher TMB may contribute to the lower sensitivity of T11-UV to the immunomodulating effects of TGFβi compared to T11-Apobec.

The use of TGFβi in combination with our high-dose PTX in POx micelles may also be a “double-punch” immunotherapeutic intervention against TNBC. PTX, in addition to its antimicrotubule activity, is also a Toll-like receptor 4 (TLR4) agonist and was shown to activate pro-inflammatory responses in macrophages [57]. TLR-4 gene is expressed in all three tumor models studied in the present work (**Supplementary Fig. S5**). Moreover, our prior study suggested that high-dose PTX-POx therapy can enhance immune cell death, repolarize tumor-associated macrophages to a pro-inflammatory M1-like state, and increase T-cell mediated immune responses in the tumor microenvironment in several TNBC tumors, including 4T1 and T11-Apobec [58]. This drug also produced long-term immune memory in 4T1 tumors [58]. Altogether, the activation of the TLR4 pathway by PTX in combination with TGFβ inhibition can further invigorate the immune tumor microenvironment across tumor models, which may explain the robust performance of the combination treatments. Future investigation delineating mutation signatures that favor better outcomes in response to the combination therapy of TGFβi and PTX could prove helpful in genetically defining treatment-responsive tumor subgroups.

In summary, our study suggests that combination therapy with TGFβis and PTX in high-capacity POx micelles provides a robust anti-tumor response in multiple TNBC subtype models. Therefore, we posit that the combination of TGFβi with PTX in high-capacity polymeric micelles could be a good remedy for TNBC.

## Acknowledgment

We thank C. Santos from the ASC for performing the tumor inoculation and M. Ross and P. V. Kantum from the ASC for performing the i.v. injections and o.g. TEM was performed by A. Shankar Kumbhar at the Chapel Hill Analytical and Nanofabrication Laboratory (CHANL), a member of the North Carolina Research Triangle Nanotechnology Network (RTNN), which is supported by the NSF (grant ECCS-1542015) as part of the National Nanotechnology Coordinated Infrastructure (NNCI). Research reported in this publication was supported in part by North Carolina Biotech Center Institutional Support Grant 2017-IDG-1025 and by the National Institutes of Health 1UM2AI30836-01. The content is solely the responsibility of the authors and does not necessarily represent the official views of the National Institutes of Health. The UNC Flow Cytometry Core Facility is supported in part by P30 CA016086 Cancer Center Core Support Grant to the UNC Lineberger Comprehensive Cancer Center. We thank the Small Animal Imaging Facility at the UNC Biomedical Imaging Research Center for providing the imaging service. The imaging core is supported partly by an NCI cancer core grant, P30-CA016086-40. We also thank the staff of Lineberger Comprehensive Cancer Center Pathology Services Core, supported by grants from the National Cancer Institute, the North Carolina Biotechnology Center (2015-IDG-1007), and the UNC Center for Environmental Health and Susceptibility and the staff of the University of North Carolina Pathology Services Core facility for providing scanning services and preparing sections for histological analysis, respectively.

We would like to acknowledge that part of this manuscript was included in the doctoral dissertation of Natasha Vinod and is publicly available in the UNC institutional library.

## Author Contributions

Background research and original draft: N.V. Review and editing: D.H., A.V.K., E.E.L., H.M.A., P.D.R., C.M.P, and N.V. Oversight: A.V.K., C.M.P., H.Y. and M.S.P. Methodology: N.V., D.H., S.C.F., T.C.O., O.C.T., S.C., J.D.R., H.M.A., E.E.L., P.D.R., and A.E. Data Analysis: N.V., D.H., P.D.R., H.M.A., E.E.L. Figure Creation: N.V., D.H., S.C.F., P.D.R., H.M.A., E.E.L.

