## Supplemental Data for "Combination of Polymeric Micelle Formulation of TGFβ Receptor Inhibitors and Paclitaxel Produce Consistent Response Across Different Mouse Models of TNBC"

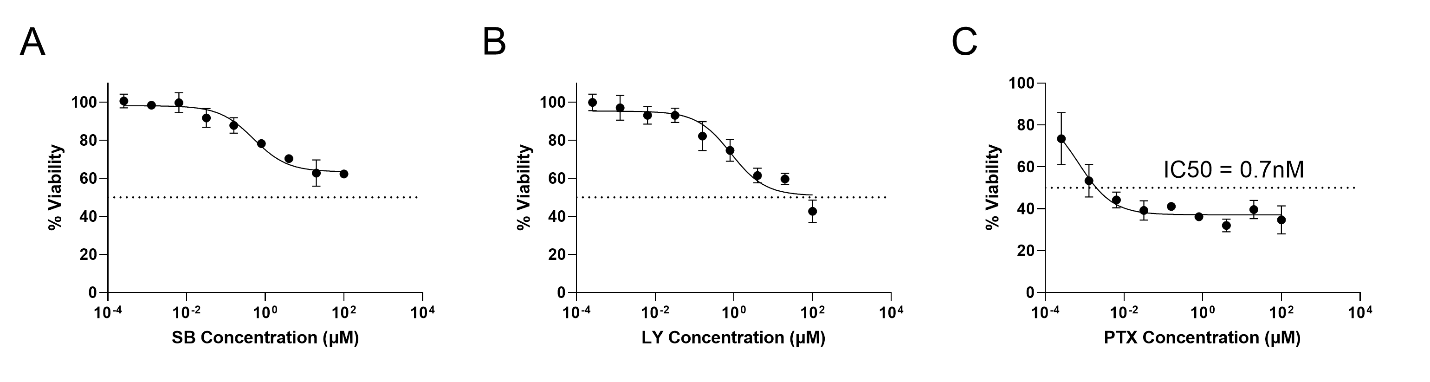
**Fig. S1 *In vitro* cytotoxicity of TGFβi and PTX in 4T1 cell line** Cell viability percent following 24h treatment with (**A**) SB-POx (**B**) LY-POx (**C**) PTX-POx.

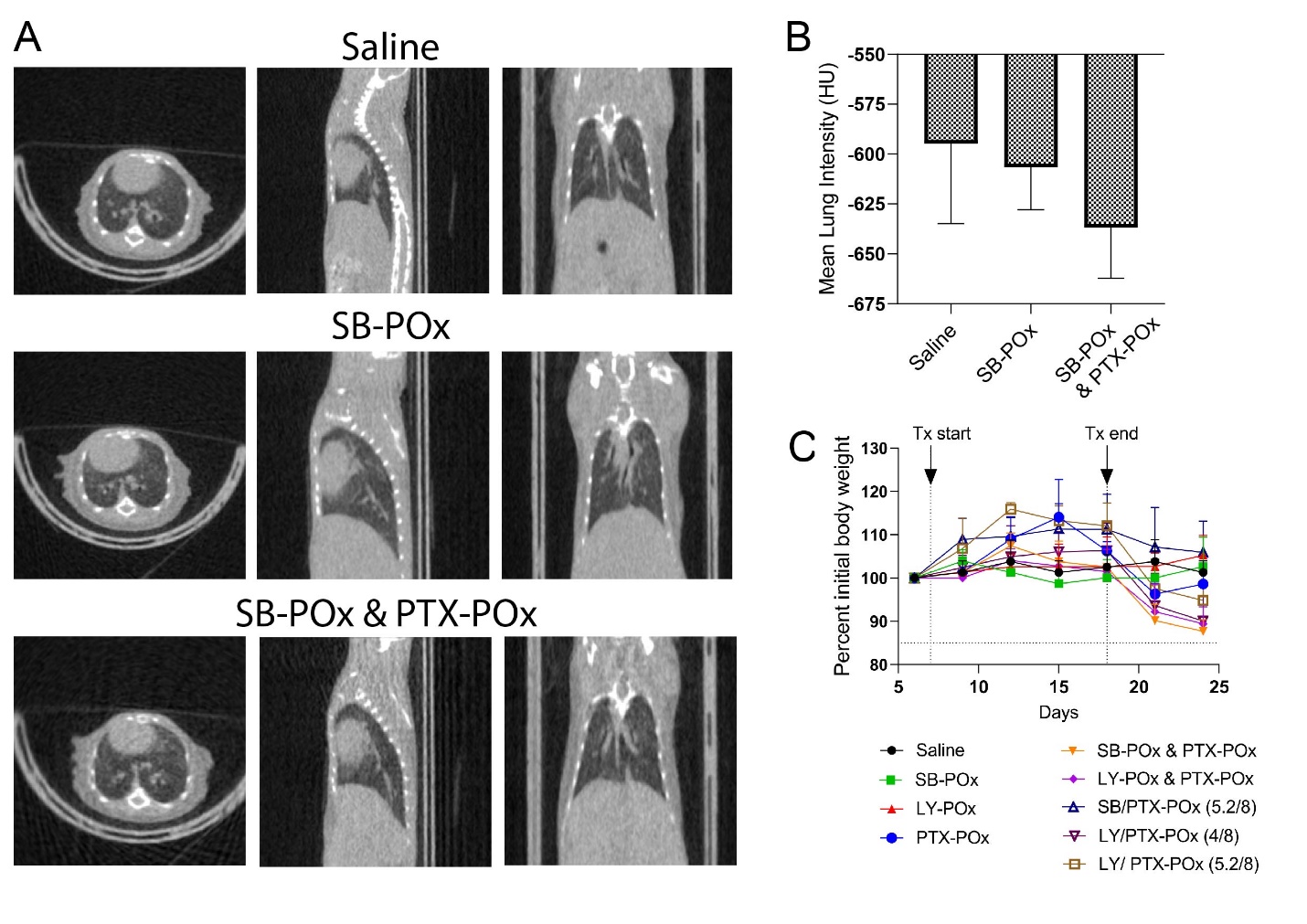

**Fig. S2** **Efficacy of intraperitoneally delivered TGFβi against primary and metastatic tumors** (**A**) Percent tumor growth inhibition corresponding to tumor growth curves in 3C and 3D (**B**) Microtomography quantification of mean lung intensity expressed in the Hounsfield units (**C**) Body weight changes (percent of initial) in mice treated with TGFβi PM and/or PTX PM.

Table S1: Experimental conditions for the preparation of POx formulations of TGFβi and PTX

| Formulation | Stock concentration of drugs in ethanol (mg/ml) | Evaporation temperature (^0^C) | Hydration temperature (^0^C) | Hydration time  (min) |
| --- | --- | --- | --- | --- |
| SB-POx | 10 | 50 | RT | 20 |
| LY-POx | 5 | 65 | Intermittent heating at 65 | 5 |
| PTX-POx | 10 | 45 | 65 | 20 |
| SB/PTX-POx (5.2/8) |  | 50 | Intermittent heating at 65 | 5 |
| LY/PTX-POx (4/8) |  | 60 | Intermittent heating at 60 | 5 |
| LY/PTX-POx (5.2/8) |  | 65 | Intermittent heating at 65 | 5 |

Table S2: Details of antibodies used in Western blot and flow cytometry experiments

| Antibody | Manufacturer | Clone | Host | Antibody Dilution | Lysate concentration |
| --- | --- | --- | --- | --- | --- |
| β-actin | Cell Signaling Technology | 13E5 | Rabbit | 1:50 | 0.4 mg/mL |
| p-Smad2/3(pThr^8^) | Sigma-Aldrich | Polyclonal | Rabbit | 1:100 | 0.4 mg/mL |
| PE anti-p-Smad2 (pS465/pS467) /Smad3(pS423/pS425) | BD Biosciences | 072-670 | Human | 1:20 | - |

**Table S3** 4T1 model. Statistical analyses of data corresponding to Figure 3A. By one-way ANOVA with Tukey’s test for multiple comparisons. Statistical difference: **** (p < 0.0001). (Graphpad Prism, version 8.4.3).

|  | PTX-POx (8/20) | SB-POx (8/20)  daily | SB-POx (8/20)  eod | SB-POx (8/20) daily + PTX-POx (8/20) | SB-POx (8/20) eod + PTX-POx (8/20) |
| --- | --- | --- | --- | --- | --- |
| Saline | **** | ns | ns | **** | **** |
| PTX-POx (8/20) |  | **** | **** | ns | ns |
| SB-POx (8/20) daily |  |  | ns | **** | **** |
| SB-POx (8/20) eod |  |  |  | **** | **** |
| SB-POx (8/20) daily  & PTX-POx (8/20) |  |  |  |  | ns |

**Table S4** 4T1 model. Statistical analyses of data corresponding to Figures 4A and 4B. By one-way ANOVA with Tukey’s test for multiple comparisons. Statistical difference: * (p < 0.05), ** (p < 0.01), *** (p < 0.001), and **** (p < 0.0001). (Graphpad Prism, version 8.4.3).

|  | PTX PM | SB PM | LY PM | SB PM + PTX PM | LY PM + PTX PM | SB/PTX (5.2/8) PM | LY/PTX (4/8) PM | LY/PTX (5.2/8) PM |
| --- | --- | --- | --- | --- | --- | --- | --- | --- |
| Saline | **** | ** | ns | **** | **** | ns | **** | **** |
| PTX-POx |  | ns | ns | ns | ns | ns | ns | ns |
| SB-POx |  |  | ns | ns | * | ns | ns | ns |
| LY-POx |  |  |  | **** | **** | ns | ** | ns |
| SB-POx & PTX-POx |  |  |  |  | ns | *** | ns | ns |
| LY-POx & PTX PM |  |  |  |  |  | **** | ns | * |
| SB/PTX (5.2/8) PM |  |  |  |  |  |  | * | ns |
| LY/PTX (4/8) PM |  |  |  |  |  |  |  | ns |

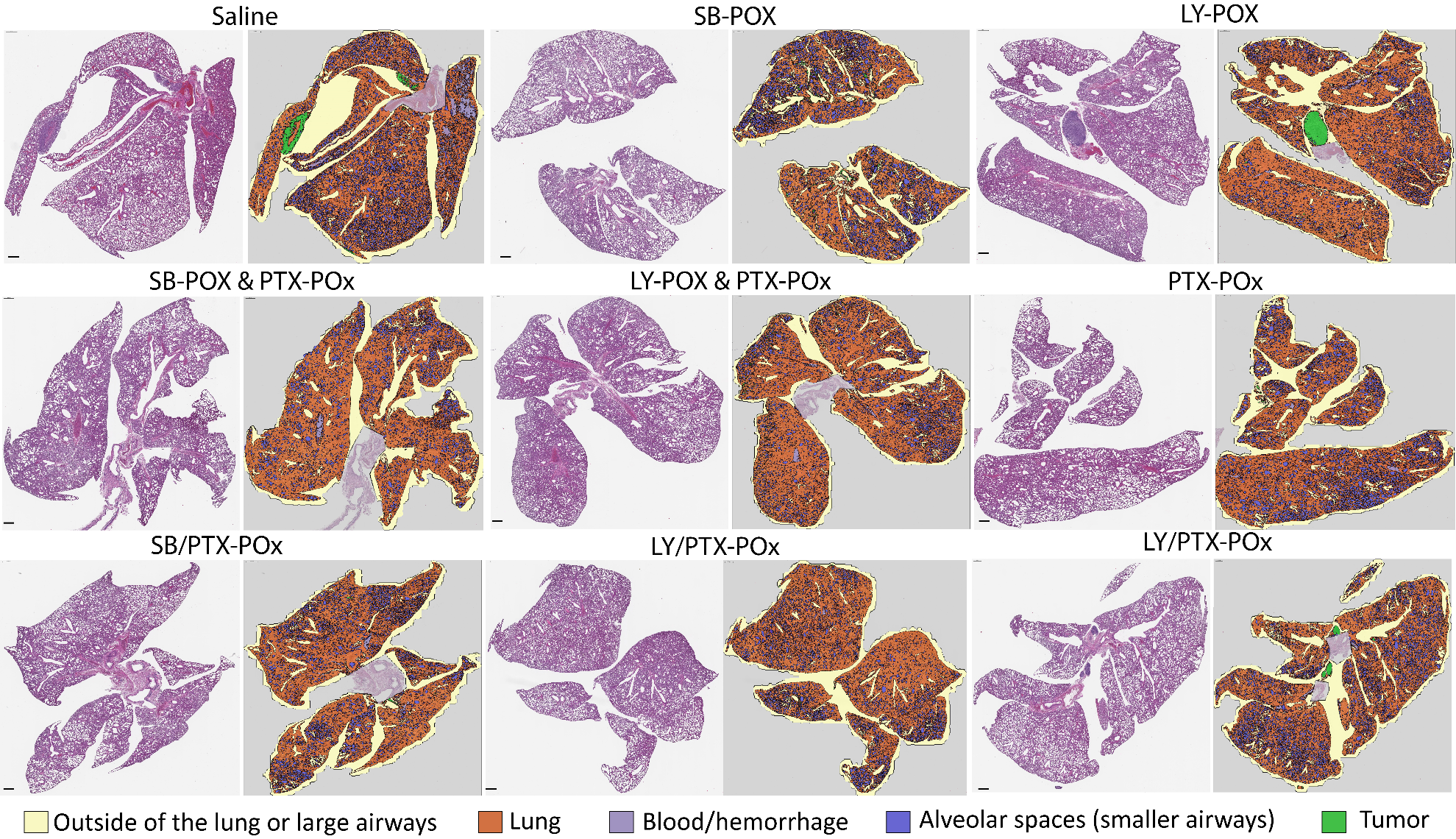

**Fig. S3 Micrometastases in the lungs of mice treated with PTX-POx and/or intraperitoneal TGFβi-Px.** Representative images of the H&E staining of mouse lungs (left) corresponding to Figure 4D with the image analysis software overlay delineating the tumor margins in green (right).

**Table S5** 4T1 model. Statistical analyses of data corresponding to Figure 5A. By one-way ANOVA with Tukey’s test for multiple comparisons. Statistical difference: * (p < 0.05), ** (p < 0.01), *** (p < 0.001), and **** (p < 0.0001). (Graphpad Prism, version 8.4.3).

|  | PTX-POx | o.g. SB- POx | o.g. LY-POx | o.g. SB-POx  & PTX-POx | o.g. LY-POx  & PTX-POx | o.g. SB | o.g LY | o.g. SB  & PTX-POx | o.g. LY  & PTX-POx |
| --- | --- | --- | --- | --- | --- | --- | --- | --- | --- |
| Saline | ** | ** | ns | **** | ns | ns | ns | **** | **** |
| PTX PM |  | **** | ** | ns | ns | *** | * | ns | ns |
| o.g. SB PM |  |  | ** | **** | **** | * | *** | **** | **** |
| o.g. LY PM |  |  |  | **** | ns | ns | ns | **** | **** |
| o.g. SB PM & PTX PM |  |  |  |  | ** | **** | **** | ns | ns |
| o.g. LY PM & PTX PM |  |  |  |  |  | * | ns | *** | ns |
| o.g. SB |  |  |  |  |  |  | ns | **** | **** |
| o.g. LY |  |  |  |  |  |  |  | **** | *** |
| o.g. SB & PTX PM |  |  |  |  |  |  |  |  | ns |

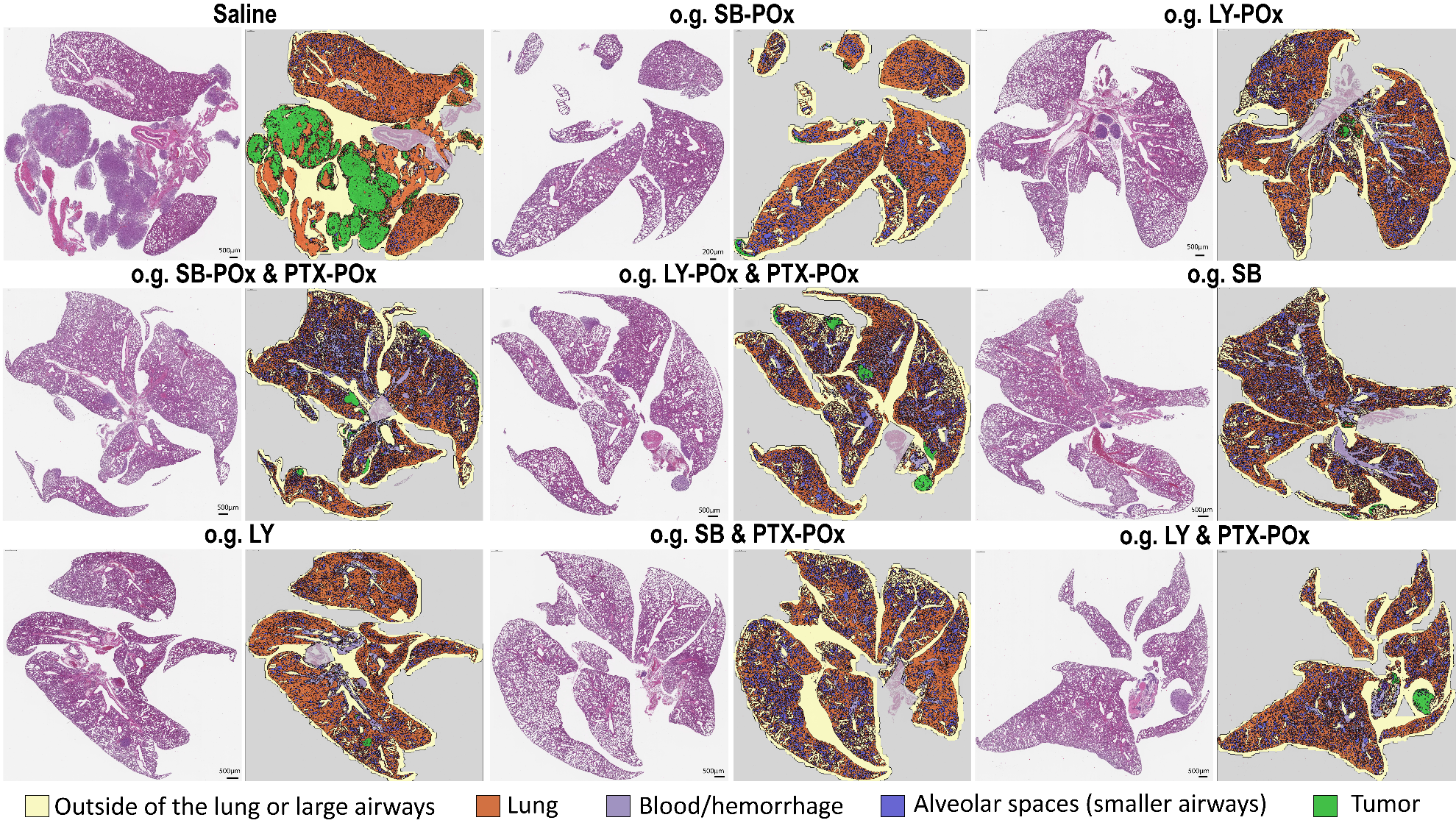

**Fig. S4 Micrometastases in the lungs of mice treated with PTX-POx and/or oral TGFβi.** Representative images of the H&E staining of mouse lungs (left) corresponding to Figure 5C with the image analysis software overlay delineating the tumor margins in green (right).

**Table S6** T11-Apobec model. Statistical analyses of data corresponding to Figure 6A. By one-way ANOVA with Tukey’s test for multiple comparisons. Statistical difference: * (p < 0.05), ** (p < 0.01), *** (p < 0.001), and **** (p < 0.0001). (Graphpad Prism, version 8.4.3).

|  | PTX PM | SB PM | LY PM | SB PM + PTX PM | LY PM + PTX PM |
| --- | --- | --- | --- | --- | --- |
| Saline | **** | **** | **** | **** | **** |
| PTX PM |  | ns | ns | * | *** |
| SB PM |  |  | ns | ns | * |
| LY PM |  |  |  | ns | ns |
| SB PM + PTX PM |  |  |  |  | ns |

**Table S7** T11-UV model. Statistical analyses of data corresponding to Figure 6B. By one-way ANOVA with Tukey’s test for multiple comparisons. Statistical difference: * (p < 0.05), ** (p < 0.01), *** (p < 0.001), and **** (p < 0.0001). (Graphpad Prism, version 8.4.3).

|  | PTX PM | SB PM | LY PM | SB PM + PTX PM | LY PM + PTX PM |
| --- | --- | --- | --- | --- | --- |
| Saline | **** | ns | ** | **** | **** |
| PTX PM |  | **** | *** | ns | ns |
| SB PM |  |  | ns | * | *** |
| LY PM |  |  |  | ns | ** |
| SB PM + PTX PM |  |  |  |  | ns |

A

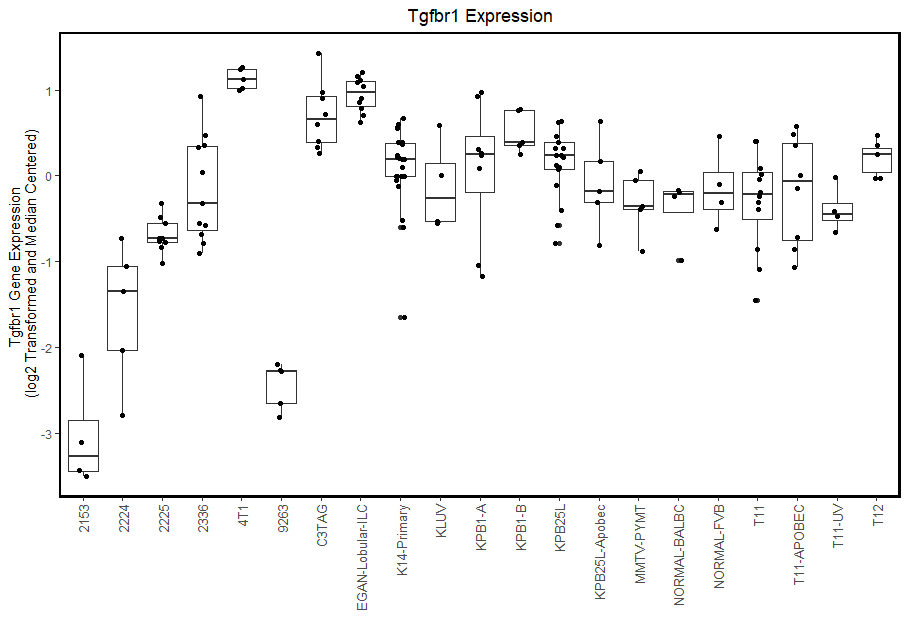

B

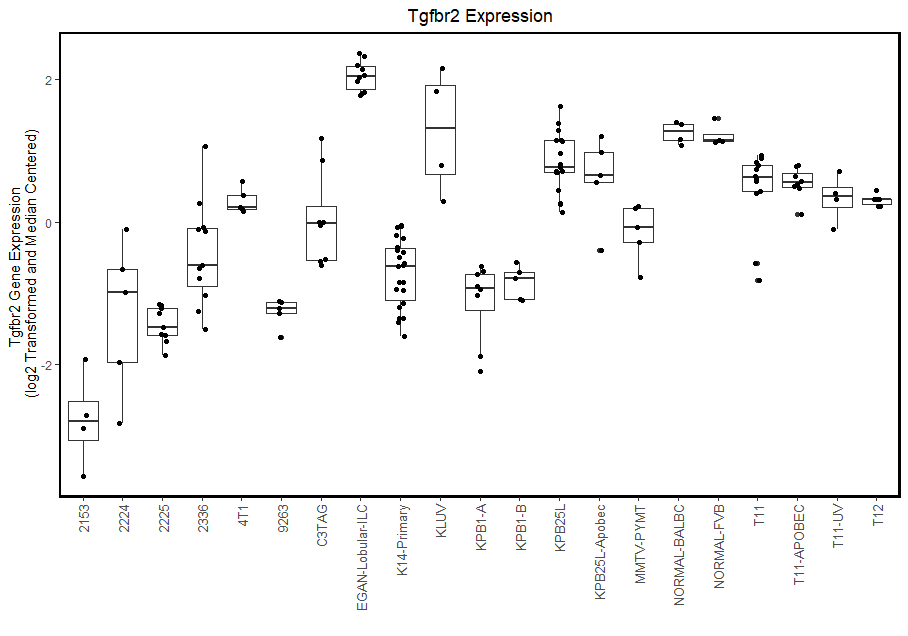

C

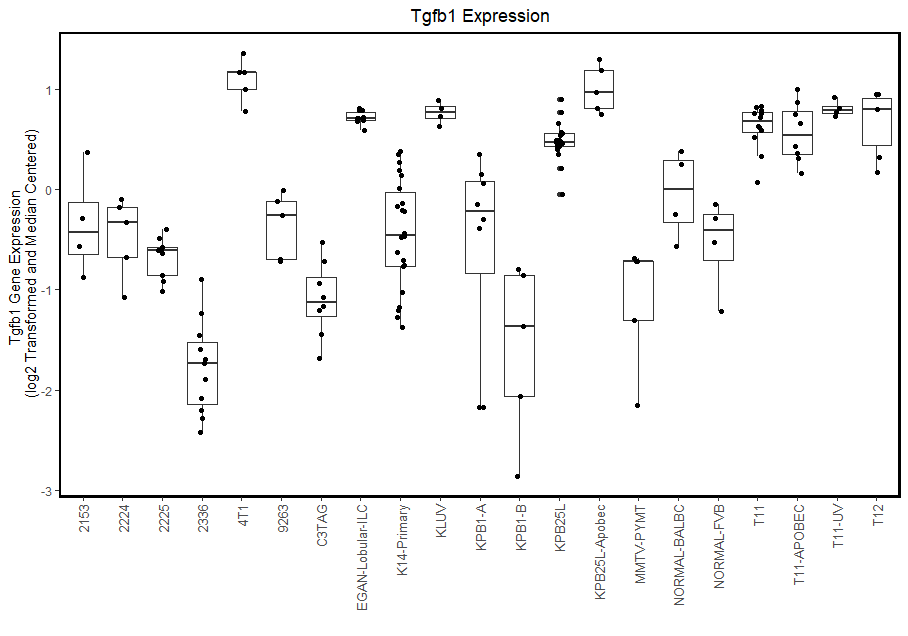

D

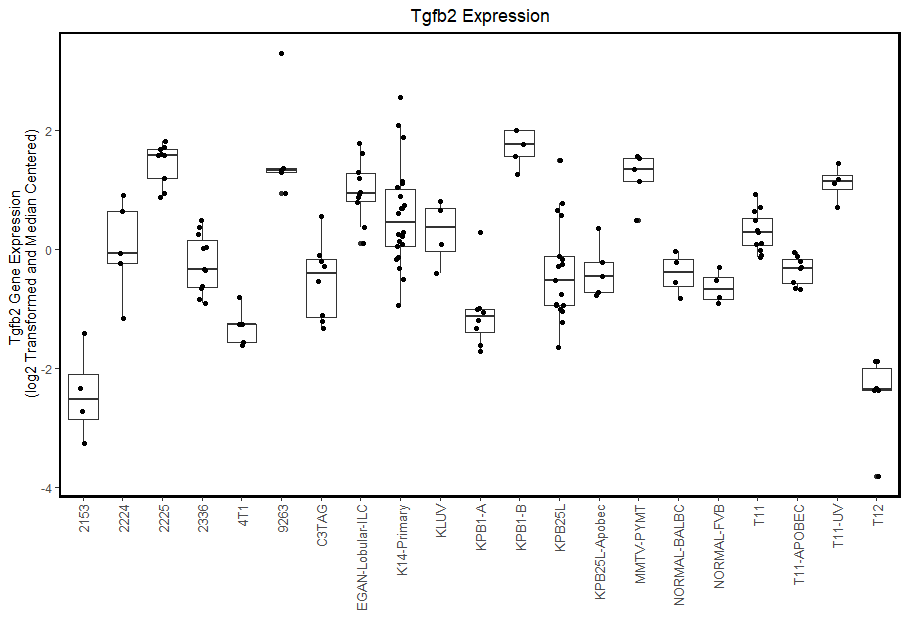

E

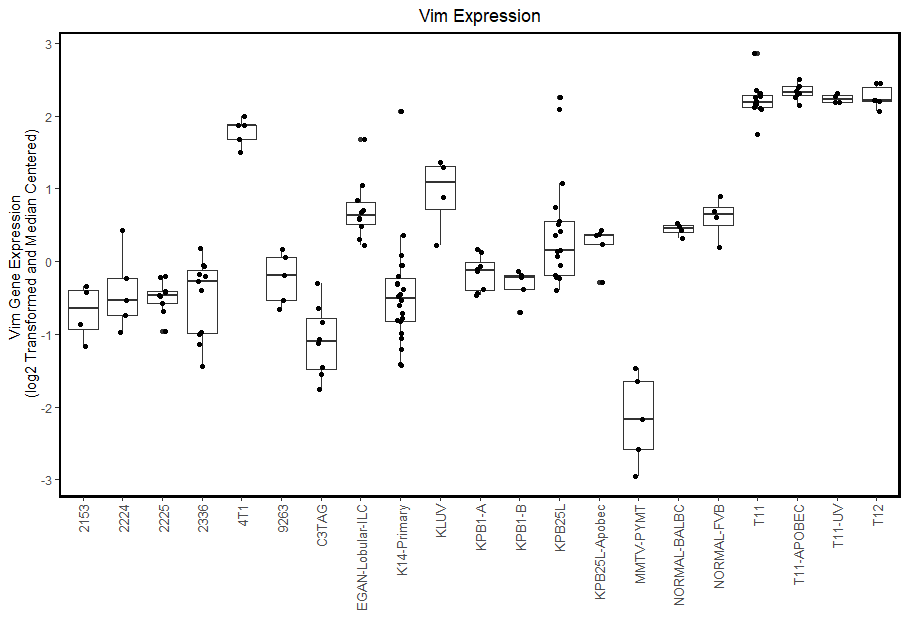

F

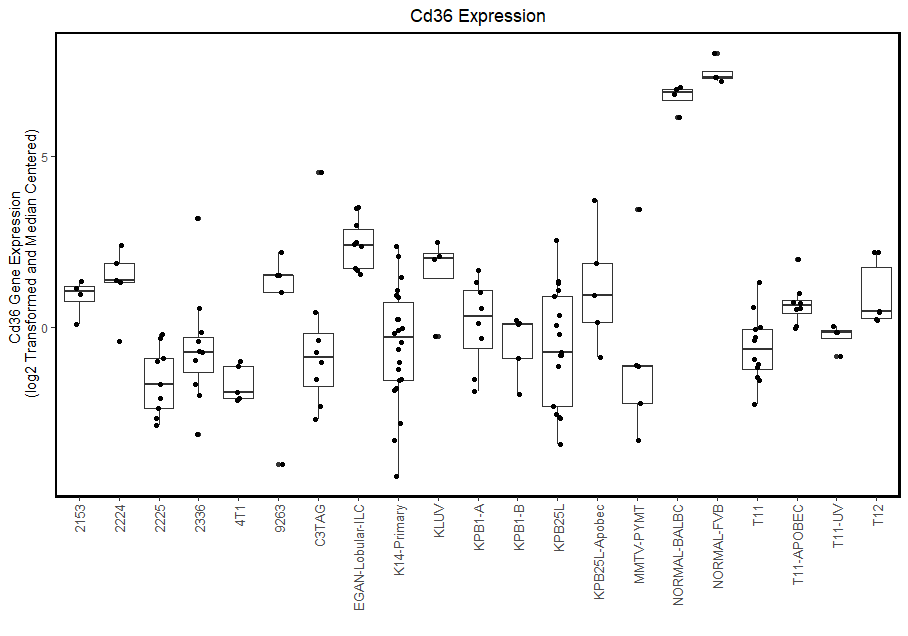

G

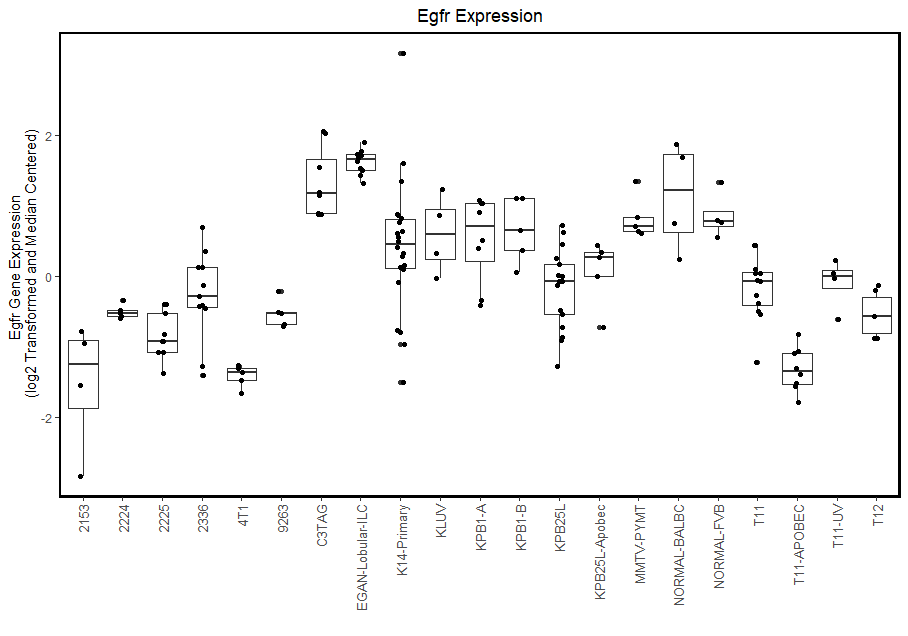

H

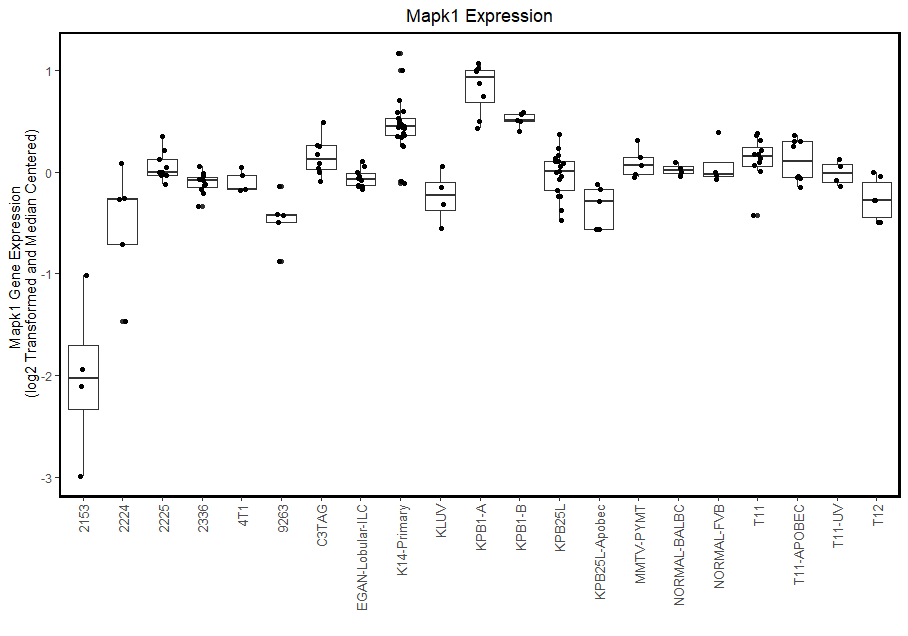

I

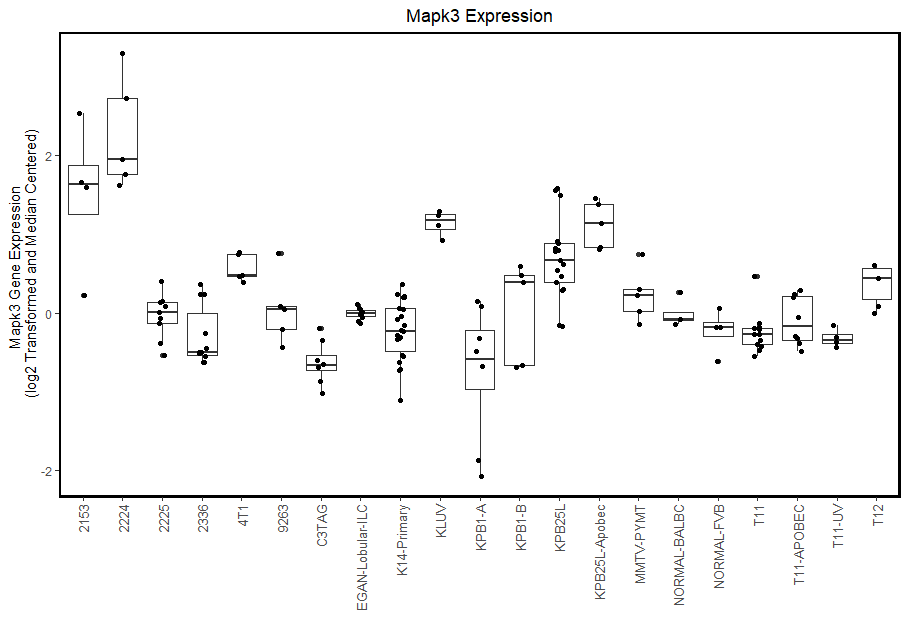

J

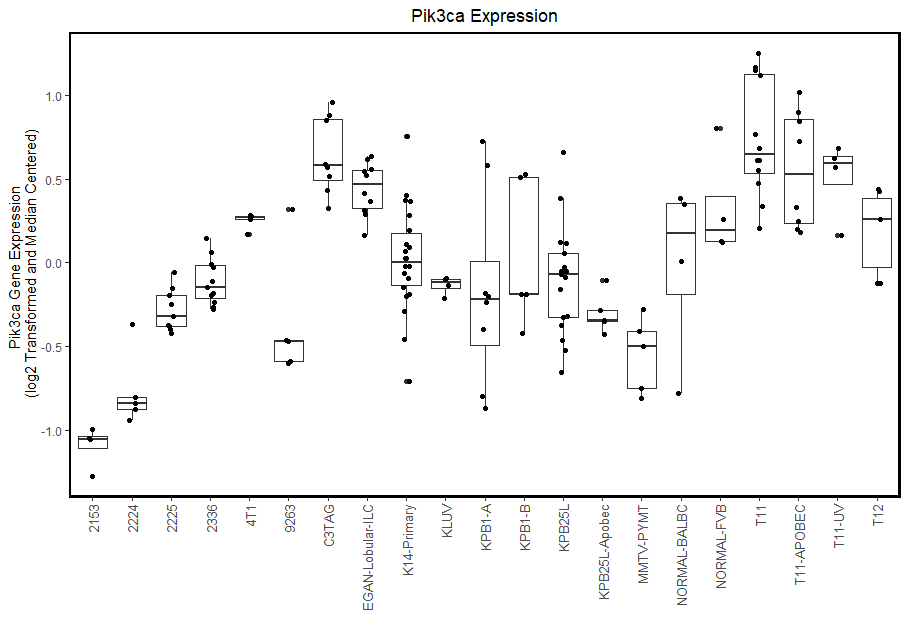

K

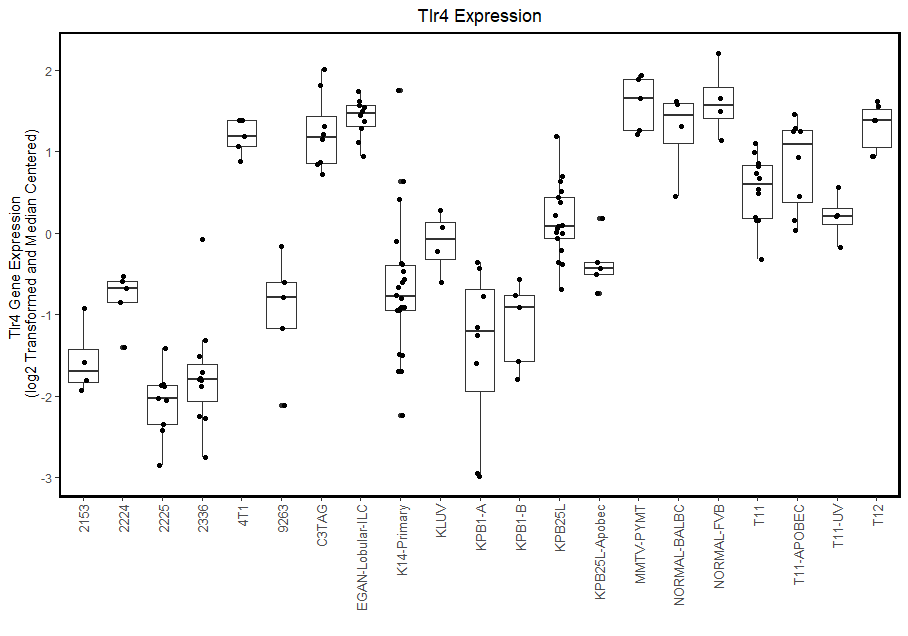

L

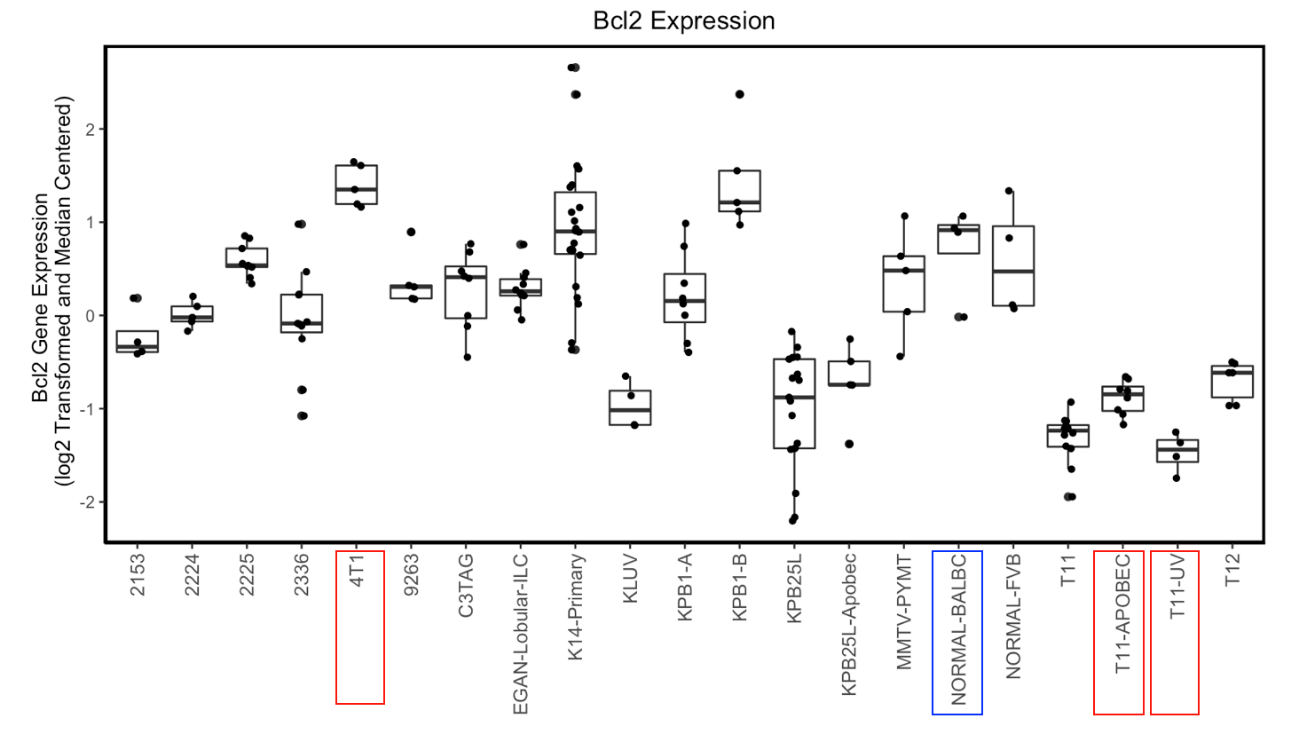

**Fig. S5.** Transcriptome evaluation (RNA-seq) of 21 TNBC models including normal breast gene expression profiles. A – transforming growth factor beta receptor 1 (Tgfbr1), B – transforming growth factor beta receptor 2 (Tgfbr2), C – transforming growth factor beta 1 (Tgfb1), D – transforming growth factor beta 2 (Tgfbr2), E – vimentin (Vim), F – CD36, G – Epidermal growth factor receptor (Egfr), H – Mitogen-Activated Protein Kinase 1 (Mapk1), I – Mitogen-Activated Protein Kinase 3 (Mapk3), J – Phosphatidylinositol-4,5-Bisphosphate 3-Kinase Catalytic Subunit Alpha (Pik3ca), K – toll-like receptor 4 (Tlr4), L – B-cell lymphoma 2 (Bcl2). The whole exome sequencing of mouse tumor was carried out as previously described (Hollern 2019).
